# A molecularly distinct cell type in the midbrain regulates intermale aggression behaviors in mice

**DOI:** 10.1101/2023.10.19.562724

**Authors:** Chunyang Li, Cheng Miao, Yao Ge, Jiaxing Wu, Panpan Gao, Songlin Yin, Pei Zhang, Hongbin Yang, Bo Tian, Wenqiang Chen, Xiaoqian Chen

## Abstract

**Background:** The periaqueductal gray (PAG) is a central hub for regulation of aggression, while little is known on the circuitry and molecular mechanisms that govern this regulation. We investigate the role of a distinct cell type, *Tachykinin 2*-expressing (Tac2^+^) neurons, located in the dorsomedial PAG (dmPAG), in modulating aggression in mice.

**Methods:** We combined activity mapping, *in vivo* Ca^2+^ recording, chemogenetic and pharmacological manipulation, and a viral-based translating ribosome affinity purification (TRAP) profiling using a mouse resident-intruder model.

**Results:** We reveal that the dmPAG^Tac2^ neurons were selectively activated during fighting behaviors. Activation of the dmPAG^Tac2^ neurons evoked, while inhibition or genetic ablation of the dmPAG^Tac2^ neurons suppressed fighting behaviors. TRAP profiling of dmPAG^Tac2^ neurons revealed that fighting behaviors specifically induced enrichment of serotonin-associated transcripts to the dmPAG^Tac2^ neurons. Last, we validated these findings by selectively delivering pharmacological agent into the dmPAG and reversed the behavioral outcomes induced by chemogenetic manipulation.

**Conclusions:** We identify that the dmPAG^Tac2^ neuron can regulate mouse aggressive behavior and thus suggest a distinct molecular target for the treatment of exacerbated aggressive behaviors in populations that exhibit high-level of violence.

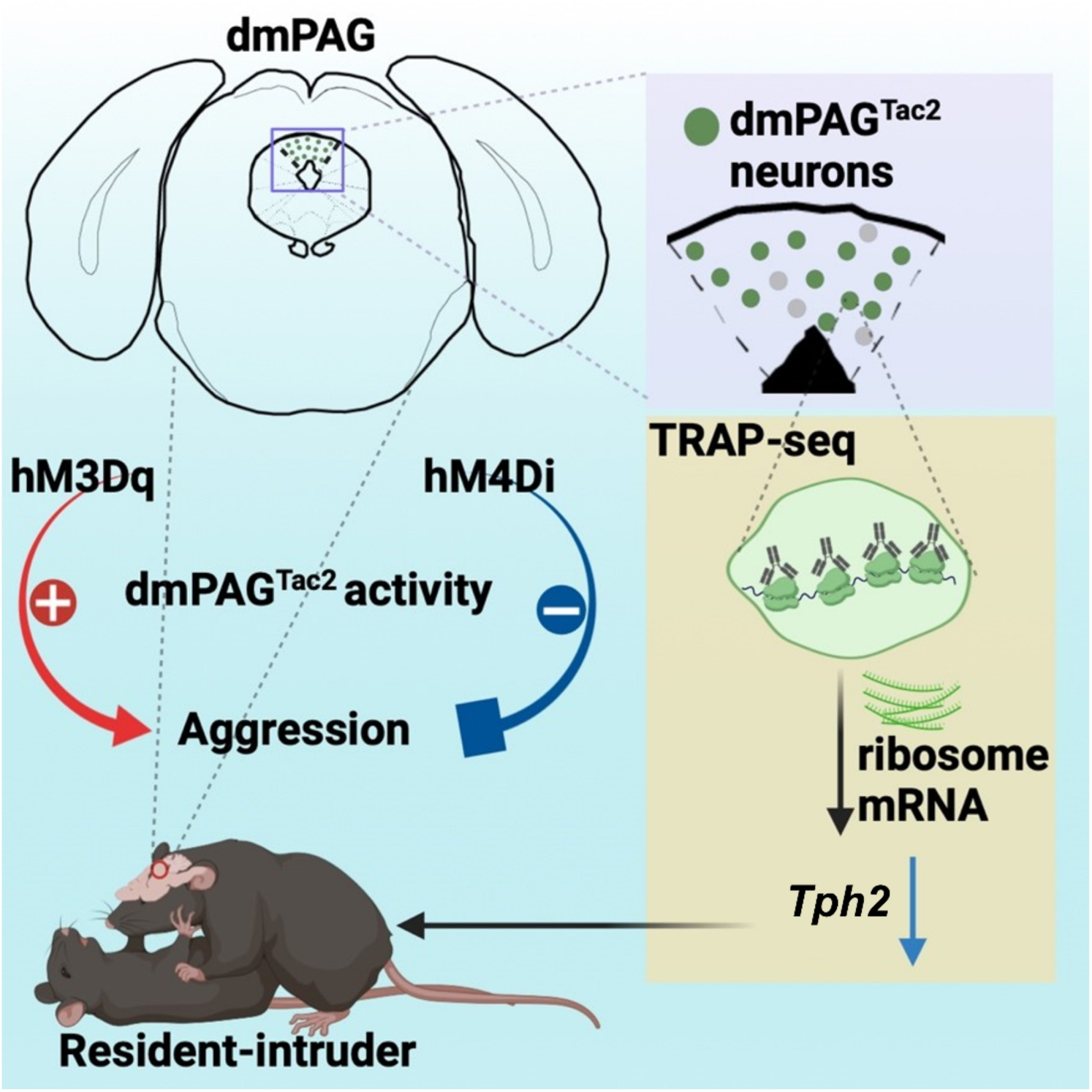

## Introduction

The periaqueductal gray (PAG) is a conserved brain region located in the midbrain traditionally viewed as a key integration center for processing upstream information related to emotion [1–3]. Early studies have demonstrated that electrical stimulation of the PAG in cats directly elicited an aggressive-defensive response, while lesions of the PAG blocked this response [4, 5]. Electrical stimulation of the amygdala and the hypothalamus also induced aggressive responses, however, when these regions are lesioned, PAG stimulation still elicited robust aversive responses [6], suggesting that the PAG may play an independent role in eliciting aversive responses such as aggression.

Functional activity-dependent studies also confirmed the critical roles of the PAG in regulating aggression and defense behaviors. More recently, studies using optogenetics, Ca^2+^ imaging, and electrophysiological recording of the PAG have revealed the causal roles of the PAG in aggressive and defensive behaviors. Of note, many of these studies were conducted in subregions such as the dorsal PAG (dPAG) and the lateral PAG (lPAG) in a pan-neuronal manner [7, 8]. However, it is unclear whether there is a distinct cell type in the PAG regulates aggressive behaviors.

In recent years, the crucial role of Tac2 positive (Tac2^+^) neurons in emotional behavior regulation has been demonstrated, including fear memory and social behaviors [9, 10]. However, most of these studies focused on Tac2^+^ neurons in limbic systems such as the amygdala and the hypothalamus, however, the critical roles of Tac2^+^ neurons in other brain regions, including the PAG, is not fully investigated. In this study, we focused on Tac2^+^ neurons in the midbrain, specifically in the dorsomedial part of the PAG (dmPAG). Using a resident-intruder (RI) paradigm in mice, we revealed a critical role of dmPAG^Tac2^ neurons in intermale aggression. Further, we explore the possible molecular targets in dmPAG^Tac2^ neurons through TRAP-seq profiling and pharmacological agent-based validation. Collectively, these findings show that the *Tac2*-expressing neurons serve as a molecularly distinct cell type in the PAG that regulates intermale aggression and thus suggesting that targeting these cells might help alleviate symptoms in humans exhibiting high levels of aggression and violence.

## Methods

### Animals

Adult (8-10 weeks) *Tac2*-Cre (Jackson, strain name B6.129-Tac2tm1.1(cre)Qima/J, stock number 018938), Ai9-tdTomato reporter mice (Jackson, stain name B6.Cg-Gt(ROSA)26Sortm9(CAG-tdTomato)Hze/J, stock number 7909), and Wild-type (WT) mice were used in this study. *Tac2*-Cre mice were kindly provided by Prof. Minmin Luo at the Chinese Institute for Brain Research, Beijing, China. Ai9-tdTomato reporter mice were kindly provided by Prof. Jian-Zhi Wang at Huazhong University of Science and Technology (HUST). All the mice were on a C57BL/6J background and were under a 12h/12h light-dark cycle with food and water provided *ad libitum*. All procedures were approved by institutional guidelines and the Animal Care and Use Committee at HUST.

### Viral vectors

AAV_2/9_-hSyn-DIO-mCherry, AAV_2/5_-EF1a-DIO-DTA-mCherry were acquired from BrainVTA Co., Ltd (Wuhan, China). AAV_2/9_-hSyn-GCaMP6m, AAV_2/9_-hSyn-DIO-GCaMP6m, AAV_2/9_-hSyn-DIO-hM3Dq-mCherry, AAV_2/9_-hSyn-DIO-hM4Di-mCherry, and AAV_2/9_-hSyn-FasDO-GCaMP6m were obtained from Taitool Bioscience Co., Ltd (Shanghai, China). AAV_2/5_-FLEX-EGFPL10a was purchased from Addgene (Watertown, USA). Titers and volumes of these viral vectors were described in the corresponding method descriptions below. All viral vectors were aliquoted and stored at -80℃ until use.

### Stereotaxic surgeries and viral injections

All stereotaxic surgeries were performed under general anesthesia using a stereotaxic apparatus (Stoelting, Wood Dale, USA). Mice were intraperitoneally anaesthetized with chloral hydrate (350 mg/kg) and xylazine (10 mg/kg) before fixed at a mouse adaptor (RWD life science, Shenzhen, China). In addition, a heating pad was also attached to the mouse adaptor to maintain a stable body temperature. After fixation on the adaptor, mice were eye-protected with ointment and transferred to a stereotactic apparatus. After the skull was exposed and leveled on the stereotactic, drilling was performed, followed by an injection of concentrated viral vector into the dmPAG (AP: -4.48 mm, ML: -0.03 mm, DV: -2.0 mm) using calibrated glass microelectrodes connected to an infusion pump (SYS-Micro4, WPI, UK) at a speed of 30 nl/min. The injection needle was held in position for 10 min and then was withdrawn. Subsequently, the incision was closed with suture. Mice were kept on a heating pad until fully recovered from anesthesia.

### In vivo fiber photometry recording

*In vivo* fiber photometry recording was performed as previously described [11, 12]. AAV-hSyn-GCaMP6m (1.1×10^13^ vg/ml, 400nl) or AAV-hSyn-DIO-GCaMP6m (2.4×10^13^ vg/ml, 200nl) was injected into dmPAG. Two weeks later, a unilateral implantation of optical fiber (200 µm, NA = 0.37; Inper LLC, Hangzhou, China) was performed on mice, which was about 200 μm above the dmPAG (AP: -4.48 mm, ML: -0.03 mm, DV: -1.8 mm). Mice were allowed to recovery at least 7 days after optical fiber implantation before the RI test. During the 15 min test, behavioral performance and calcium activity of dmPAG neurons were recorded simultaneously. Specifically, fiber-photometry system consisted of a 473-nm LED excitation light redirected by a dichroic mirror featuring a reflection band of 435–488 nm and a transmission band of 502–730 nm. The light was then focused into a 200 μm 0.37 NA optical fiber using an objective lens. To investigate dmPAG^Tac2^ neuronal activity under various stimuli, we recorded GCaMP6s fluorescence intensity in mice during the RI test, object exploration, exposure to fox odor, or pain stimulation, respectively. Raw data of Ca^2+^ fluorescence (F) were collected by fiber photometry (sampling at a frequency of 25 Hz) and normalized by baseline correction, motion correction and signaling filtering using Inper Data Process Software (Inper LLC). The Ca^2+^ fluorescence was then aligned with the onset of corresponding behavioral events, and the fluorescence change values (dF/F) were calculated as (F– F0)/F0, where F0 represents the baseline fluorescence signal averaged over a 4-second time-window which was from 6 seconds to 2 seconds prior to the event onset. The calculation, data process, analysis and visualization were performed by MATLAB (R2017a) and GraphPad Prism (v8).

### Behavioral assays

All mice were transferred to the testing room for the habituation of the environments at least 3 h prior to all tests. When the entire testing trial of each mouse finished, all behavioral apparatus were cleaned with 75% ethanol to eliminate the odor left by the previous mice. Mice were tested during the dark phase, with all behavioral experiments conducted in darkness.

### Open-field test (OFT)

The OFT was performed as previously reported with minor modifications [12–14]. Briefly, a white open-field box (50 cm × 50 cm × 50 cm) made of PVC materials was used. During the test, mice were placed gently in the center of the open arena as a starting position and were allowed to explore freely for 10 min. Data were recorded and analyzed using SuperMaze software (Shanghai Xinruan Informatlon Technology Co.Ltd, Shanghai, China). Total distance travelled in the open-filed arena, time spent in the central zone and total numbers of entries into the central zone were analyzed.

### Object exploration test

The object exploration was performed as previously reported with minor modifications [15]. This test was conducted in the same open-field arena for the OFT test. Briefly, after a short adaptation to an empty open-field arena, mice were placed into the open-field arena containing an inanimate object for 5 min. Behavioral performance and Ca^2+^ fluorescence of the dmPAG^Tac2^ neurons were recorded and aligned to the onset of mice exploratory behaviors, that is, when mice directed the nose toward the object at a distance less than or equal to 2 cm.

### Elevated-plus maze (EPM) test

The EPM test was performed as previously reported with minor modifications [12–14]. Briefly, we used an EPM apparatus placed 62 cm in height from the ground. This apparatus was made of two open arms (75 cm × 5 cm) and two closed arms (75 cm × 5 cm × 15 cm) (Shanghai Xinruan Informatlon Technology Co.Ltd). To start the test, mice were placed gently in the center area of the maze, with the mice facing to the open arm. Mice behaviors in the maze were recorded for 8 minutes using SuperMaze software. Time spent in open/closed arms and numbers of total entries into open/closed arms were analyzed.

### Rotarod test

To measure muscle strength and motor coordination in mice, the rotarod test was performed as previously described with minor modifications [12–14]. Briefly, a revolving cylinder (6 cm in diameter) with a rough surface to increase grasp was used. Prior to the start of the test, mice received training to adapt the rotating rod with a low speed (accelerated from 5 to 10 rpm) for 3 sessions per day. Each session lasted for 30 min and was set with at least a 30-min interval. During the test, we adjusted the speed of the rotarod accelerated from 5 to 40 rpm over a 5-min period. Each mouse was tested for four trials a day with a 30-min interval between each trial. The detector automatically recorded the moment each mouse dropped off from the revolving rod. Latency to drop was counted and analyzed.

### Three-chamber social interaction test

The three-chamber social interaction test was conducted in a rectangular plexiglass box (60 cm × 40 cm × 22 cm) as reported previously [12, 13]. The testing box was separated into 3 equal chambers by 2 clear plexiglass walls, each contains an entryway that mice can move with free access. A wire cage (5 cm in diameter and 10 cm in height) was placed in the center of the 2 side chambers. To start each test, mouse was placed in the middle chamber as the starting point. For session1 (habituation), empty wire cage was placed in the two side chambers, and the mouse was allowed to explore the apparatus freely for 10 min. During session2 (sociability test), a stranger social mouse, whose species and sex were the same as the testing mouse, was placed inside one of the wire cages at one side chamber, then the testing mouse was allowed to explore the apparatus freely for 10 min again. During session3 (social novelty test), a new stranger social mouse, whose species and sex were the same as the testing mouse but coming from a different cage as the previously caged mouse, was placed inside another wire cage that was empty during session2. The testing mouse was allowed to explore the apparatus freely for another 10 min. There are a 5-min inter-session interval after each session. Data were recorded, retrieved and analyzed by using SuperMaze software. Social preference score was calculated as (T_S_−T_NS_)/(T_S_+T_NS_), in which T_S_ or T_NS_ represented the time that a testing mouse spent in the social area or non-social area, respectively.

## 2-methyl-2-thiazoline (2MT) odor stimuli

2MT odor stimuli assay was conducted as previously described with minor modifications [16–18]. For fiber photometry recordings of dmPAG^Tac2^ neurons to fear stimuli of 2MT, a potent analog of fox odor 2,4,5-trimethyl-3-thiazoline (TMT) was used to elicit robust innate fear/defensive behaviors in mice, such as freezing. The mice tested were recorded in their own cage and single housed at the test room. After 5 min with control odor (saline) exposure, 20 μl of 2MT (2.1 × 10^−4^ mole) was added on filter paper in a small dish and place it away from the mice and in the corner farthest from them, test and record for 5 min again.

### Hot plate pain test

Hot plate pain test was performed as previously described [16, 19]. Mice were placed on a 275 mm × 263 mm × 15 mm metal plate and were subjected to a 5-min acclimation at 30°C firstly, then the temperature increased or decreased at the rate of 6℃/min which the maximum setting at 55°C until the mice manifested aversive behaviors, including hind-paw licking, shaking, lifting or jumping, video of nociceptive heat response was recorded.

### Resident-intruder (RI) test

RI test is a classical paradigm to test aggressive behaviors in mice [20] while its operational details vary depending on research focus and purpose [21]. Here, “resident” mice subjected to RI test were housed with a female companion for at least 7 d before the test, which increased their sense of territory. During this period, the same bedding material was kept clean. We did not choose the commonly aggression-increasing paradigm, in which the “resident” mice were single-housed several days prior to the RI test, as previous study has reported that chronic social isolation broadly upregulated Tac2/NkB signaling across the brain areas [10]. One hour before RI test, the female companion mouse was removed from the cage, and a stranger C57/bl6 male “intruder” mouse, which was slightly smaller in body size than the resident, was introduced into the cage and allowed to interact for 15 min freely. An effective attack was defined as the resident mouse exhibited biting, wrestling, tumbling towards the conspecific during the RI test [22–24]. Other behavioral events during the RI test, such as sniffing, defined as a nose contact made by the resident to any part of the intruder’s body including genital sniff [25], and social grooming [26] were also counted and analyzed. Videos were recorded, and latency to attack, numbers of attack episode, and duration of attack were analyzed using Adobe Premiere Pro CC 2018 and GraphPad Prism 8.0.

### Chemogenetic modulation of dmPAG^Tac2^ neurons

Chemogenetic manipulation stimulated by Clozapine-N-oxide (CNO) (intraperitoneal injection) was performed as previously reported with minor modifications [12–14]. According to literatures [11, 24, 25, 27, 28], CNO dosage can be varied ranging from 0.1 to 0.7 mg/kg for chemogenetic aggression activation and from 0.5 to 7.5 mg/kg for chemogenetic aggression inhibition. Based on our piloting tests, we chose CNO dosage (1mg/ml, C0832, Sigma-Aldrich) of 0.5 mg/kg for chemogenetic aggression activation and 1.0 mg/kg for chemogenetic aggression inhibition.

For chemogenetic activation of dmPAG^Tac2^ neurons, *Tac2*-Cre mice was microinjected with AAV-hSyn-DIO-hM3Dq-mChery (Gq, 1.22×10^13^ vg/ml, 200 nl) into the dmPAG (*Tac2*^hM3Dq^ mice). Control *Tac2*^mCherry^ mice received AAV-hSyn-DIO-mChery (5×10^12^ vg/ml, 500 nl). According to a previous report [25] on potential “ceiling effect”, all mice underwent two rounds of preselection before virus injection to select mice with low baseline levels of spontaneous attacks. Twenty-one days after virus injections, all mice were tested for aggression for 3 consecutive days to confirm their low spontaneous attacks. Then mice received either CNO (0.5 mg/kg) or normal saline (control) and were used for RI test 30 min later. Using normal home-cage housing *Tac2*^hM3Dq^ mice, the efficacy of CNO-activated Gq virus in dmPAG^Tac2^ neurons was verified by c-Fos staining at 120 min after CNO administration.

For chemogenetic inhibition of dmPAG^Tac2^ neurons, *Tac2*-Cre mice with high baseline levels of spontaneous attacks were selected before virus injection to avoid potential “flooring effect” [25]. Eligible mice received either AAV2-hSyn-DIO-hM4Di-mChery (Gi virus, 1.77×10^13^ vg/ml, 150 nl, *Tac2*^hM4Di^) or AAV2-hSyn-DIO-mChery (*Tac2*^mCherry^). Twenty-one days after injection, mice were tested for their higher spontaneous attacks for 3 consecutive days. The *Tac2*^hM4Di^ or *Tac2*^mCherry^ mice were then administrated with either CNO (1.0 mg/kg) or normal saline for RI test.

### Genetic ablation of dmPAG^Tac2^ neurons

*Tac2*-Cre mice with high baseline levels of spontaneous attacks were selected for genetic ablation assay. To genetically ablate dmPAG^Tac2^ neurons, AAV-hSyn-DIO-DTA-mCherry (5.89×10^12^ vg/ml, 450 nl) was injected into the dmPAG of *Tac2*-Cre mice. *Tac2*-Cre mice receiving AAV-hSyn-DIO-mCherry virus were served as control. Twenty-one days following viral injection, mice were subjected for RI test.

### Immunofluorescence

Immunofluorescent staining was performed as reported previously [29, 30]. Briefly, mice were perfused with ice cold normal saline, followed by perfusion with ice code 4% paraformaldehyde. After perfusion, whole brains were carefully removed from the skull and fixed in 4% paraformaldehyde for 6 h at 4°C for post-fixation and then transferred to 20% and 30% sucrose solution for gradient dehydration.

To explore and verify expression of Tac2 across the whole brain, brains from *Tac2*-Cre/Ai9-tdT double transgenetic mice were used (tdTomato^+^ in Tac2^+^ neurons). After gradient dehydration, coronal sections were prepared from frozen brain tissues of *Tac2*-Cre/Ai9-tdT mice via Leica 1860 vibratome at a thickness of 30 μm. Brain slices were rinsed in phosphate-buffered saline (PBS) 3 times (5 min each) and then were stained with Hoechst (1:1000, #33342, Sigma-Aldrich) for 10 min. These slices were rinsed in PBS 3 times and were cover-slipped with fluorescent mounting medium and proceeded with photography immediately.

To verify immunofluorescent expression and injection site of virus, brains of mice that were subjected to stereotaxic surgeries were obtained. Coronal frozen sections of the PAG region were prepared at a thickness of 20 μm. Sectioned brain slices were rinsed in PBS 3 times and then stained with Hoechst for 10 min. These slices were rinsed in PBS 3 times and were cover-slipped with fluorescent mounting medium. After staining, slices were proceeded with photography immediately.

For c-Fos staining, we followed a previous report [31]. Brains of mice that were subjected to resident-intruder test were sacrificed 90 min after the test or sacrificed 120 min after receiving CNO administration. Coronal frozen sections of the whole-brain areas (for c-Fos mapping) or the PAG area (for CNO administration) were prepared at a thickness of 20 μm. Subsequently, the brain slices were permeabilized with 0.3% Triton X-100 in PBS for 10 min and blocked with 5% goat serum in PBS for 1 h. The brain slices were then incubated with primary antibodies (1:800, rabbit monoclonal anti-cFos antibody, #2250S, Cell Signaling) at 4°C overnight. Following incubation with primary antibodies, brain slices were rinsed in PBS 3 times and then were incubated in fluorochrome-conjugated secondary antibody (1:200, Dylight^488^-conjugated goat anti-rabbit, Abbkine; 1:200, AF^488^-conjugated goat anti–rabbit IgG, Jackson ImmunoResearch) for 1 h at room temperature in dark. These slices were then rinsed in PBS 3 times and then stained nuclei with Hoechst for 10 min. Afterwards, the slices were rinsed in PBS 3 times again, cover-slipped with fluorescent mounting medium and stored in 4°C prior to photography.

### Translating ribosome affinity purification (TRAP) and RNA sequencing

Affinity purification of translating ribosomes was performed as described with minor modifications [32]. Briefly, *Tac2*-Cre mice were subjected to stereotaxic viral injection of AAV-EF1a-FLEX-EGFPL10a (7×10^12^ vg/ml, 400 nl) at dmPAG. Three weeks later, brain areas containing dmPAG were rapidly dissected (individual dmPAG were pooled from 6-7 mice per sample; thus, each group there were 3 replicates) at 4°C (cold room) and were immediately transferred into ice-cold dissection buffer for tissue wash to remove blood. These samples were then transferred to a pre-chilled homogenizer on ice that contains tissue-lysis buffer (20 mM HEPES, pH 7.4, 10 mM MgCl_2_, 150 mM KCl, 0.5 mM DTT, 100 mg/ml cycloheximide, 10 ml/ml RNasin and Superasin). We used a set of pre-cooled douncer-based method which containing a loose pestle douncer and a tight pestle douncer (Kimble douncer for 20 times. Homogenates were then centrifuged at 2,000g, 10 min, 4°C, to pellet large cell debris. NP-40 (Beyotime Biotechnology, Shanghai) and 1,2-diheptanoyl-sn-glycero-3-phosphocholine (Avanti Polar Lipids, AL) were added to the supernatant at final concentrations of 1% and 30 mM, respectively. After incubation on ice for 5 min, the lysates were centrifuged for 10 min at 20,000g to pellet insoluble materials. A small amount of supernatant (about 5%) was collected in separated centrifuge tubes, which representing “*Input*” fraction. GFP antibodies (Proteintech, 5 μg) were added into the remaining supernatant for an end-over-end incubation (4 h, at 4°C) on a rotator. Protein A+G magnetic beads (Beyotime Biotechnology) were prepared during the incubation. The cleaned beads were then added to the tissue homogenates when the incubation was finished. The tissue homogenates were allowed to continue the end-over-end incubation for another 16-18 h at 4°C. The next day, when the incubation was finished, the beads were resuspended with high-salt buffer (20 mM HEPES KOH, pH 7.3, 350 mM KCl, 10 mM MgCl_2_, and 1% NP-40, 0.5 mM DTT, 100 μg/ml cycloheximide and Rnasin) at least 4 times. After the last cleaning of the beads, pre-cooled centrifuge tubes were placed on the magnet holder for at least 1 min, and all remaining cleaning fluid was carefully removed. Afterwards, RLT buffer (Qiagen Kit 74004, DTT added) were added to the tubes, and these tubes were removed from the magnet holder and vortexed for 30 s at room temperature. Subsequently, the tubes were placed on the magnet holder for at least 1 min at room temperature. Then, the RLT buffer that containing RNAs was transferred to new enzyme-free centrifuge tubes, which representing “*IP*” fraction. To isolate RNAs from “*Input*” fractions, RLT buffer was added to the *Input* samples and vortexed thoroughly. RNA extraction was performed side-by-side with the *IP* samples according to the instructions of RNA isolation kits. Finally, extracted RNAs or RLT lysate containing RNAs was sent to Shanghai Bohao Biotechnology for RNA quality inspection and subsequent library construction and mRNA constant sequencing analysis.

### Intra-dmPAG drug delivery for pharmacological validation

Pharmacological validation experiments were based on previous reports with minor modifications [9, 10, 33–37]. First, AAV2-hSyn-DIO-hM3Dq-mCherry was injected into dmPAG of *Tac2*-Cre mice. Two weeks later, these mice were implanted with a guide cannula at dmPAG (AP: -4.48 mm, ML: -0.03 mm). The depth of the injection catheter base was determined according to the final injection site in the target brain area and the depth of the matching injection tube. For instance, the injection site of the dmPAG of 25 g mice in this study was -2.0 mm. If the implantation depth of the injection catheter base was -1.40 mm, an injection tube of G1 = 0.6 mm was used. The implantation depth of the drug delivery catheter in each mouse is recorded so that the corresponding length of the injection tube can be used in the subsequent drug administration. The catheter cap should be inserted into the implanted drug delivery catheter to maintain the patency of the catheter, to prevent thrombosis, and to reduce intracranial infection when the drug is not injected daily.

Seven days after cannula implantation, fluoxetine (MCE, 6 mg/500nl) or vehicle (saline with DMSO) were delivered into dmPAG. Prior to behavioral tests, the guide cannula (RWD Life Science, Shenzhen, China) was removed from the testing mice and replaced with injection cannula, which protruded 0.5 - 0.7 mm from the tip of the guide cannula according to the implantation sites of the guide base. The Injection cannula was attached to a 10-ml Hamilton syringe with PE tubing. Testing drugs were delivered through the injection pump at a rate of 0.25 ml/min to inject a total volume of 500 nl of drug. When the drug delivery was finished, the injection cannula was held in place for another 3 min to allow fully diffusing the drugs into the dmPAG and to prevent the backflow of the testing drugs while the needle pulled out. Afterwards, the injection tube was carefully and slowly pulled out, and the remaining drugs in the PE tube were continued to push out to verify the patency of the needle. After intracranial injection, CNO was intraperitoneally administered immediately. The guide cannula was sterilized, and then inserted into the guide base. Behavioral tests were performed around 30 min after the CNO administration.

### Quantification and statistical analysis

The data obtained were processed and analyzed by various softwares when necessary (Excel, GraphPad, ImageJ). Statistical analyses used were unpaired *t*-test, paired *t*-test, one-way analysis of variance (one-way ANOVA), two-way ANOVA for the homogeneity of variance of data with normal distribution. Tukey test was used for multi-comparisons between specific groups following ANOVA. Data are expressed as mean ± standard error (mean ± s.e.m.). Statistical difference was indicated with * *P* < 0.05, ** *P* < 0.01, *** *P* < 0.001, **** *P* < 0.0001. Specifically, one-way ANOVA followed by Tukey’s multiple comparisons test analysis was used to compare fluorescent staining in different brain region, paired *t*-test was used to compare behavioral data of chemogenetic manipulation and intra-dmPAG drug microinfusion as previously reported [11], unpaired *t*-test was used to compare the behavioral results of WT mice and *Tac2*-Cre mice in open-field test, elevated plus maze test, and rotarod test, and two-way ANOVA followed by Tukey’s multiple comparisons test was used to analyze behavioral data of three-chamber social interaction test, repeated measures two-way ANOVA was used to compare behavioral data of DTA ablation of dmPAG^Tac2^. In addition, behavior data from resident-intruder test were collected by Xinsoft XR-XZ301 video acquisition and analysis system, and video analysis and statistics were performed by Premiere Pro CC 2018. A double-blind analysis of these videos was performed by at least two people not aware of the group information, and ultimately the mean of the two statistically collected data was taken for further statistical analysis.

## Results

### *Tac2*-expressing neurons exhibit distinct expression pattern in the PAG

Previous studies reported that Tac2^+^ neurons are expressed in a wide range of brain areas including the amygdala, hypothalamus, nucleus accumbens [38], however, their expression pattern in the PAG has not been characterized. To this end, we crossed *Tac2*-Cre knock-in mice with Ai9 reporter mice to selectively express tdTomato fluorescent protein in Tac2^+^ cells (**Figure 1A**). By aligning expression of Tac2^tdTomato^ cells to the Allen Brain Atlas (ABA), we confirmed that Tac2^tdTomato^ cells were located in selected brain areas such as the hypothalamus and amygdala (**Figure 1B-C**), consistent with previous reports [10, 38]. No tdTomato^+^ cells were found in control mice including the *Tac2*-Cre mice, Ai9 mice, and wide-type mice (**Figure S1A**).

**Figure 1.**
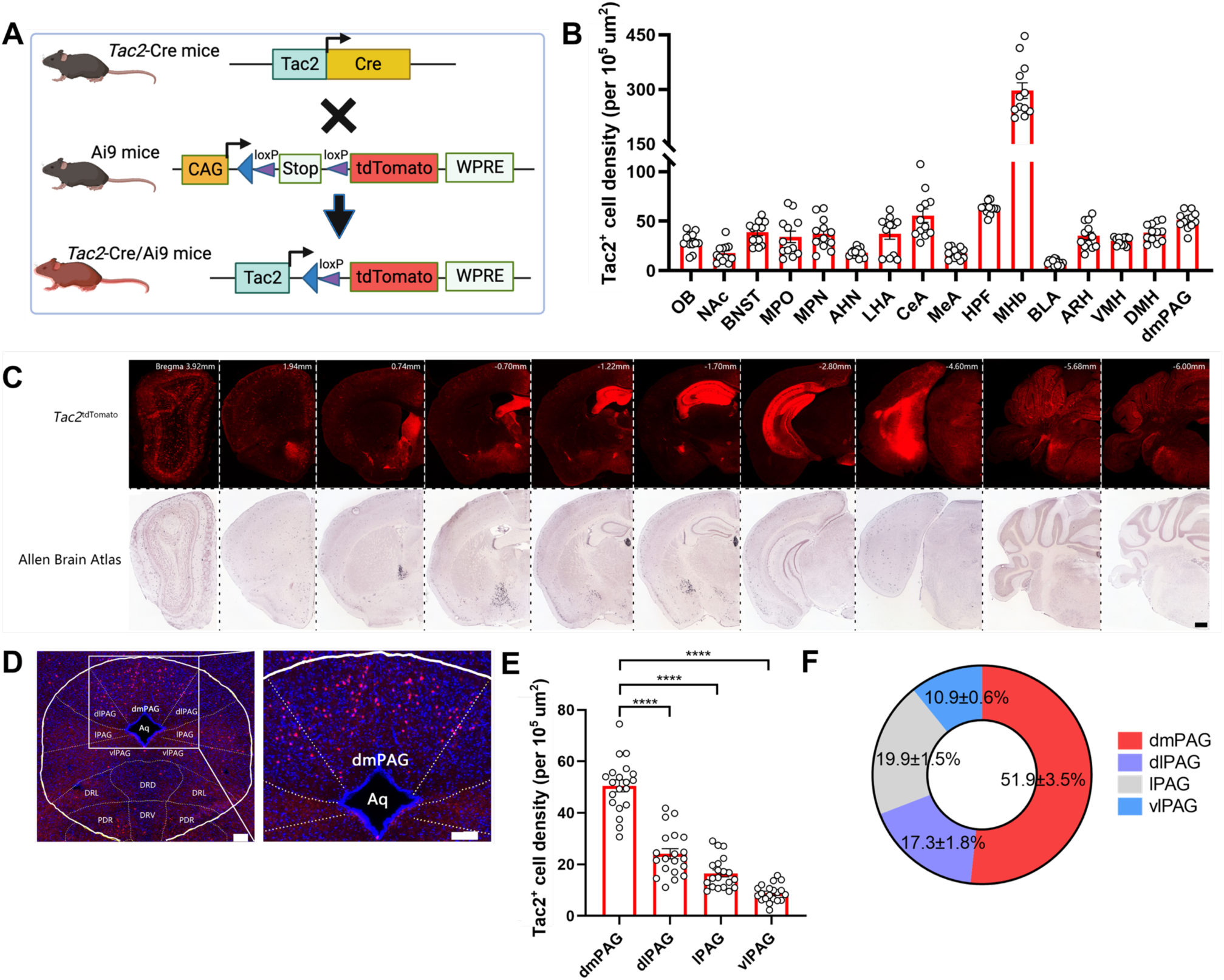
PAG Tac2-expressing neurons are mainly located in the dmPAG. **(A)** Schematic showing breeding strategy to selectively express tdTomato reporter in Tac2^+^ cells. **(B)** Whole-brain analysis of *Tac2*^tdTomato^ cell density. N=3 mice, with 4 brain sections from each mouse. **(C)** Representative fluorescent images showing tdTomato-labeled Tac2^+^ cells (red) in coronal brain sections in *Tac2*-Cre/Ai9 mice (upper panels) and in their respective ABA images (lower panels). Scale bar, 500 μm. **(D)** Distribution pattern of Tac2^+^ cells in the PAG subregions. Scale bars, 1,000 μm (left) and 100 μm (right, enlarged view), respectively. **(E)** Analysis of *Tac2*^tdTomato^ cells in PAG subregions. N=5 mice, with 4 brain sections from each mouse. **(F)** Pie chart showing percentage of *Tac2*^tdTomato^ cells in PAG subregions. A total of 648 Tac2^+^ cells were included from N = 3 mice. Data were expressed as mean ± S.E.M. Significance was calculated by One-way ANOVA and Tukey’s multiple comparisons test, **** P < 0.0001.

We then specifically analyzed the expression pattern in the PAG. Tac2^tdTomato^ cells were found throughout the PAG (**Figure 1D**), aligning with the mRNA expression pattern observed in the ABA (**Figure S1B**). Notably, Tac2^tdTomato^ cells were predominantly located in the dmPAG (**Figure 1E-F**), comprising about 50% of the total *Tac2*-expressing cells in the pan-PAG areas. This distribution is consistent with the enriched presence of *Tac2* mRNAs, which was predominantly expressed in the dmPAG (54.4%, **Figure S1C-D**). Thus, these data confirm that *Tac2*-expressing cells in the PAG are primarily located in the dorsomedial subregion.

### dmPAG^Tac2^ neurons respond to aggression

We next sought to determine the involvement of the dmPAG^Tac2^ neurons in fighting behaviors. First, we subjected the WT mice to RI assay and harvested brain tissues after fighting (**Figure 2A**). As expected, fighting behaviors activated neurons in a broad range of brain areas, including the paraventricular nucleus of the thalamus, periventricular zone, anterior hypothalamic nucleus (**Figure S2**). Focusing on the c-Fos^+^ neurons in the dmPAG, our results showed that, compared to the control mice kept at home cages, a significant number of c-Fos^+^ neurons were found in the dmPAG of the resident mice (**Figure 2B-C**). In contrast, fighting behaviors had minimal effects in inducing c-Fos expression in other PAG subregions, such as the dlPAG, lPAG, and vlPAG (**Figure S2B-C**).

**Figure 2.**
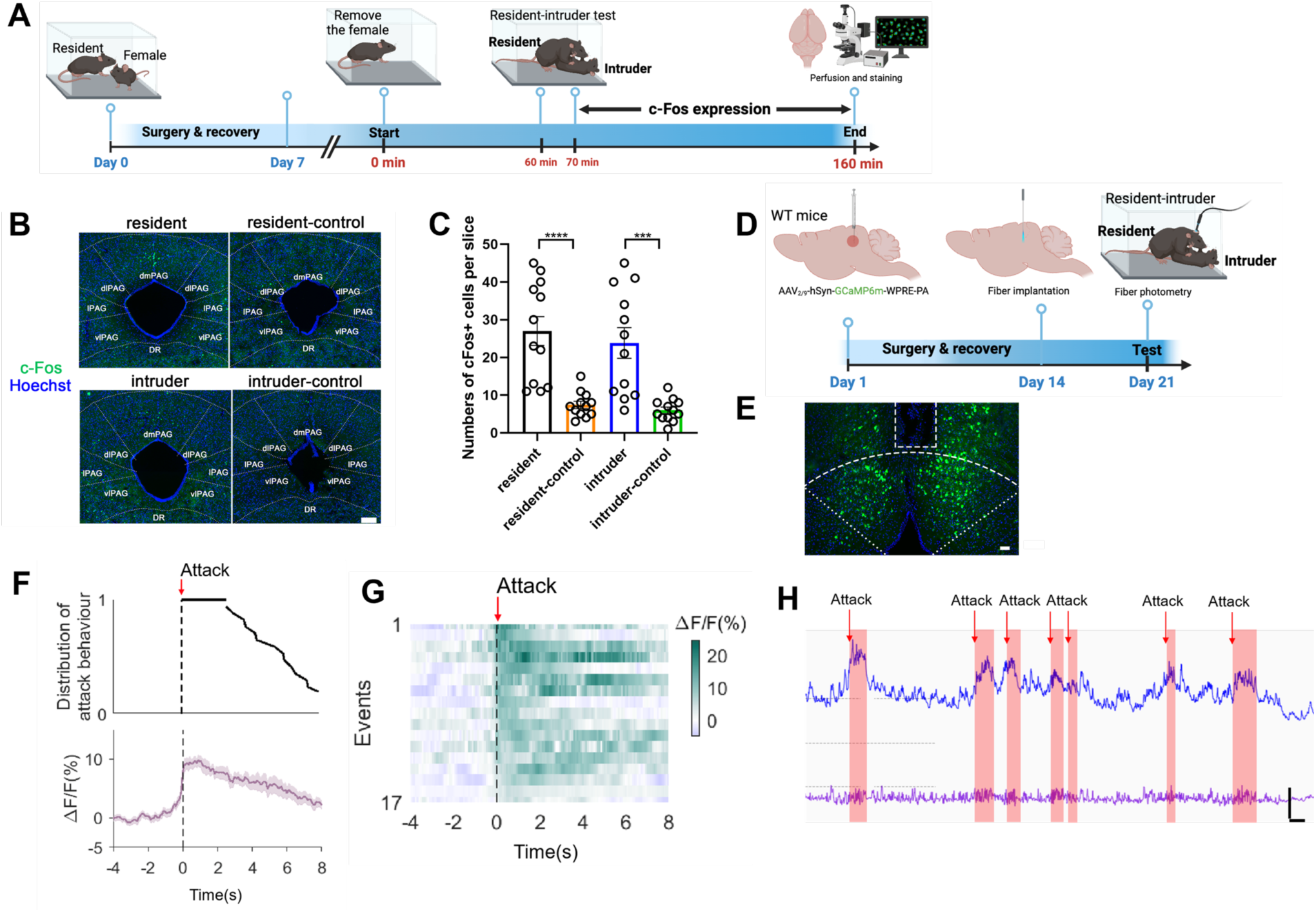
dmPAG neurons respond to aggression. **(A)** Strategy for mapping c-Fos expression in WT mice subjected to the RI test. See methods for detailed procedure. **(B)** c-Fos activation in dmPAG of mice from various groups. Scale bar, 100 μm. **(C)** Number of c-Fos^+^ cells in the dmPAG of WT mice. N = 3 mice/group, with 4 brain sections from each mouse. **(D)** Strategy for in vivo fiber photometry in WT mice injected with AAV-hSyn-GCaMP6m. **(E)** Fiber position targeting the dmPAG of WT mice. Scale bar, 50 μm. **(F)** Distribution of attack behavior synchronized with ΔF/F%. **(G)** Heatmap showing Ca^2+^ activity time-locked to an attack episode. x-axis shows 4 sec prior to and 8 sec after the attack episode (red arrow). **(H)** Representative Ca^2+^ trace showing one trial of Ca^2+^ activity in a WT mouse during the RI test. Scale bar: 10s in x-axis and 2000 A.U. in y-axis. Baseline (bottom trace): reference control channel (405 nm); upper trace, Ca^2+^ channel (470 nm). Red shaded areas indicate attack episodes and durations. Data represent mean ± S.E.M. Significance was calculated by One-way ANOVA and Tukey’s multiple comparisons test, *** P < 0.001, **** P < 0.0001.

We then investigated the role of dmPAG neurons in response to fighting behaviors using *in vivo* fiber photometry. AAV expressing GCaMP6m was injected into the dmPAG of WT mice and then a fiber was implanted above the dmPAG (**Figure 2D**). Correct placement of the fiber was verified (**Figure 2E**). When subjecting these mice to RI test, we observed a large increase in Ca^2+^ transients in response to each attack bouts (**Figure 2F-G**). Notably, each attack elicited an increase in Ca^2+^ activity immediately after the start of the attack episodes and this increase returned to baseline level after an attack (**Figure 2H**). Thus, these results demonstrated the involvement of dmPAG neurons in fighting behaviors.

We further investigated the role of dmPAG^Tac2^ neurons in fighting behaviors. We used a Cre-dependent AAV vector (**Figure 3A**) to label dmPAG^Tac2^ neurons and then subjected the mice to the RI (**Figure 3B**). Staining c-Fos proteins in mCherry^+^ cells confirmed that dmPAG^Tac2^ neurons were activated by fighting behaviors (**Figure 3C**). We then measured Ca^2+^ activity of dmPAG^Tac2^ neurons during the RI by injecting AAV carrying a Cre-dependent GCaMP6m vector (**Figure 3D-E**) and verified the correct placement of the optic fiber (**Figure 3F**). We observed an increase in Ca^2+^ activity in response to attack episode (**Figure 3G-I**). When correlating each component of an attack event to Ca^2+^ activity (**Figure 3G**), we found that attacks reliably induced Ca^2+^ transient, but none of the other attack-associated events, such as tail rattling, sniffing, and grooming, could significantly alter Ca^2+^ activity (**Figure 3J-O**). In addition, using a Cre-out strategy, we revealed that dmPAG Tac2-negative neurons showed a mild but a significant increase in Ca^2+^ activity following each attack episode (*P*<0.05, **Figure S3A-D**). This increase was, however, three times lower than that of the dmPAG^Tac2^ neurons (*P*<0.0001, **Figure S3E-F**), suggesting that, while the other cell types in the dmPAG might contribute, at least partially, to the fighting behaviors, consider the degrees of increase in Ca^2+^ activity (**Figure 3I** and **Figure S3D-F**), the dmPAG^Tac2^ neurons still exhibited a dominant role in response to aggression.

**Figure 3.**
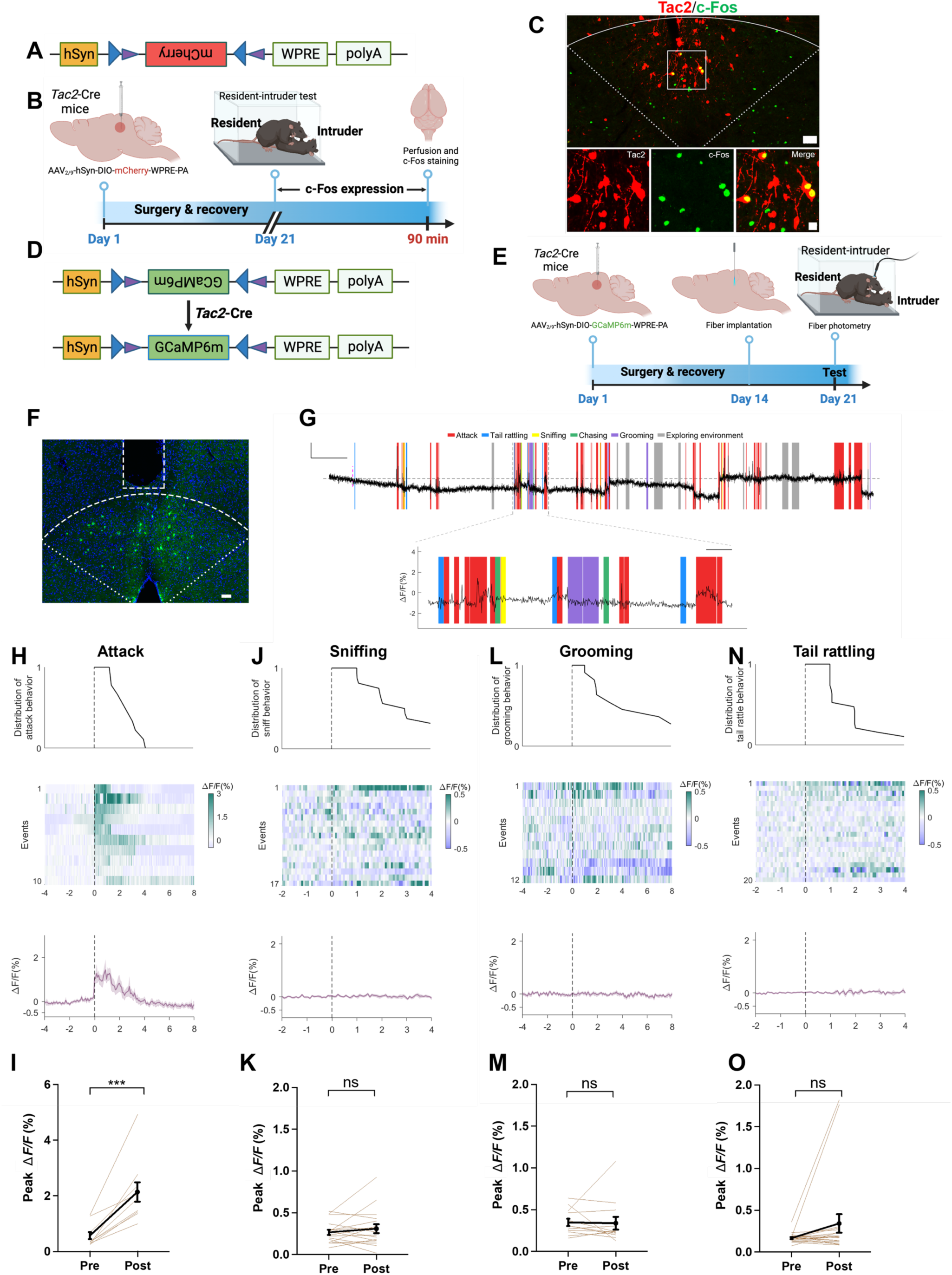
Tac2-expressing neurons in dmPAG respond to aggression. **(A)** Vector information for AAV-hSyn-DIO-mCherry. **(B)** Strategy for examining activation of Tac2^+^ neuron by fighting behaviors. AAV vector was injected to *Tac2*-(green) in dmPAG^Tac2^ neurons (red mCherry labeled) in *Tac2*-Cre mice subjected to the RI test. Lower panels showing enlarged view from the white box in the upper panel. Scale bars, 50 μm (upper), 20 μm (lower), respectively. **(D)** Vector information for AAV-hSyn-DIO-GcaMP6m in *Tac2*-Cre mice. **(E)** Strategy for in vivo fiber photometry in *Tac2*-Cre mice injected with Cre-dependent AAV-GcaMP6m vector. **(F)** Fiber position targeting dmPAG region of Tac2 mice. Scale bar, 50 μm. (G) Example recording of dmPAG^Tac2^ neurons during the RI test. Color shades indicate different behavioral syllables during intruder encounter. Scale bar: 60s in x-axis, 1.5% GCaMP6m ΔF/F signal in Y-axis. **(H-O)** Ca^2+^ activity time-locked to attack **(H-I),** sniffing **(J-K),** grooming **(L-M)** and tail rattling **(N-O)**. **(H)** Distribution of attack events (top), heatmap of GCaMP6m ΔF/F signals (middle) and ΔF/F% (bottom), **(I)** Analysis of peak ΔF/F before (Pre) and after (Post) each attack episode. The other panels present the same manner but with different behavioral syllables, including sniffing **(J-K)**, grooming **(L-M)**, and tail rattle **(N-O)**. Data represent mean ± S.E.M. Paired *t*-test, ***P < 0.001.

The PAG is a highly heterogenous region and is involved in many important functions, including sensorimotor function and pain sensation [2, 39]. To further test the specificity of dmPAG^Tac2^ neurons in fighting behaviors, we subjected these mice to a variety of stimuli and recorded Ca^2+^ activity. The results showed that, dmPAG^Tac2^ neurons failed to exhibit significant altered Ca^2+^ activity in response to male-male and male-female encounter (**Figure S4A-F**), rotarod activity (**Figure S4G-H**), object exploration (**Figure S4I-J**), fox odor sniffing (**Figure S4K-L**), saline sniffing (**Figure S4M-N**), and pain stimulation (**Figure S4O-P**). Thus, these data collectively demonstrated that, the dmPAG^Tac2^ neurons respond specifically to fighting behaviors.

### Deletion or inhibition of dmPAG^Tac2^ neurons attenuates fighting behaviors

To further confirm that dmPAG^Tac2^ neurons respond to fighting behaviors, we performed chemogenetic manipulations to examine a causal role of dmPAG^Tac2^ neurons in aggression. First, we performed a basic behavioral characterization of *Tac2*-Cre mice and their WT littermate controls. The results showed that *Tac2*-Cre mice exhibited similar levels of locomotor performances (**Figure S5A-E**). However, *Tac2*-Cre mice had mild but significantly increased levels of anxiety (**Figure S5F-J**). In addition, *Tac2*-Cre mice exhibited no difference in social preference (**Figure S5K-P**) and notably, *Tac2*-Cre mice had similar levels of fighting behaviors in the RI (**Figure S5Q-T**), which establishes a non-different baseline level of fighting behaviors prior to our next genetic ablation and chemogenetic manipulation experiments.

We then examined the behavioral effects of genetic ablation of dmPAG^Tac2^ neurons by injecting a neuron-specific Cre-dependent AAV encoding DTA-mCherry (**Figure 4A-B**) into the dmPAG of *Tac2*-Cre mice (*Tac2*^DTA^). *Tac2*-Cre mice injected with a Cre-dependent AAV-mCherry (**Figure 4A-B**) were served as the control (*Tac2*^mCherry^). DTA-mediated ablation resulted in an 80% ablation of dmPAG^Tac2^ neurons (**Figure 4C-D**), demonstrating high efficiency in ablating *Tac2*-expressing cells in the dmPAG. *Tac2*^mCherry^ mice showed similar levels of aggression before and after AAV injection, however, mice with genetic ablation of dmPAG^Tac2^ neurons exhibited a significantly decreased level of aggression (**Figure 4E-G**), when compared not only to the baseline levels (**Figure 4B**), but also to their *Tac2*^mCherry^ control mice. In addition, ablation of the dmPAG^Tac2^ neurons significantly reduced the percentage of mice that exhibited fighting behaviors (**Figure S6A**), whereas this manipulation failed to alter male-male investigation during the encounter (**Figure S6B**). Thus, these data suggest that ablation of dmPAG^Tac2^ neurons attenuates aggression behaviors in mice.

**Figure 4.**
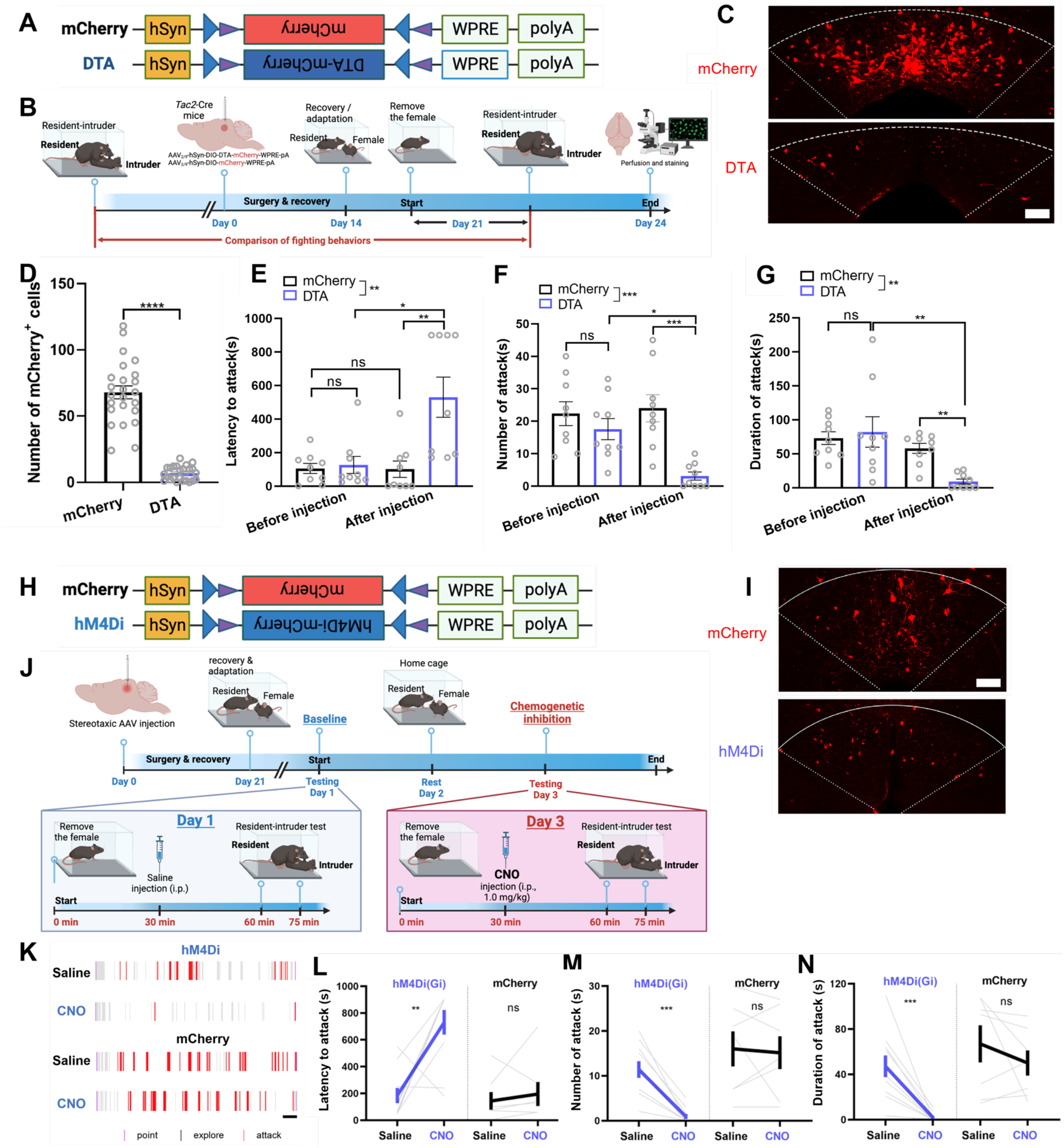
Ablation or inhibition of dmPAGTac2 neurons suppresses fighting behaviors. **(A)** A Cre-dependent DTA vector or its control vector to be expressed in dmPAG^Tac2^ neurons. **(B)** Strategy for genetic ablation. **(C)** Validation of DTA (bottom) and control AAV (upper) in *Tac2*-cre mouse. Scale bar, 100 μm. **(D)** Number of mCherry^+^ cells in dmPAG in *Tac2*-Cre mice receiving genetic ablation. N = 6 mice, with 4 brain sections from each mouse. **(E-G)** Analysis of latency to attack **(E)**, number of attack **(F)** and duration of attack **(G)**. N = 9/group. Significance was calculated with Two-way RM ANOVA followed by Tukey’s multiple comparisons test. *P < 0.05, **P < 0.01, ***P < 0.001. **(H)** A Cre-dependent hM4Di vector or its control to be expressed in dmPAG^Tac2^ neurons. **(I)** Representative fluorescent images of mCherry (upper) or hM4Di-mCherry (bottom). Scale bar, 100 μm. **(J)** Strategy for the RI test in *Tac2*-Cre mice receiving inhibitory DREADDs vectors. **(K)** Raster plots showing mouse attack (red bar) or exploring behaviors (gray bar) during the RI assay. Scale, 1 min. **(L-N)** Analysis of latency to attacks **(L)**, number of attacks **(M)** and duration of attacks **(N)**. N = 10 in *Tac2*^hM4Di^ group and N = 7 in *Tac2*^mCherry^ group. Data represent mean ± S.E.M. Significance was calculated by means of paired *t*-test. **P < 0.01,

Similarly, we performed chemogenetic silencing of dmPAG^Tac2^ neurons. First, AAV with Cre-dependent inhibitory DREADDs vector (hM4Di-mCherry) was injected into the dmPAG of *Tac2*-Cre mice (**Figure 4H-I**, *Tac2*^hM4Di^ mice). *Tac2*-Cre mice injected with a Cre-dependent AAV encoding mCherry construct were served as the control (*Tac2*^mCherry^). We established a baseline level of aggression by administering *Tac2*-Cre mice with normal saline and subjecting to the RI test after waiting for 30 min. (**Figure 4J**). Forty-eight hours later, these mice were intraperitoneally injected with CNO (1 mg/kg) and subjected to the RI again after waiting for 30 min. As expected, saline administration failed to alter fighting behaviors in *Tac2*^mCherry^ mice (**Figure 4K-N**), however, the previously established fighting behaviors in the baseline test were dramatically suppressed by CNO administration in *Tac2*^hM4Di^ mice (**Figure 4L-N**). Also, inhibition of dmPAG^Tac2^ neurons significantly reduced the percentage of mice exhibiting fighting behaviors (**Figure S6C**). Of note, we chose a relatively low dose of CNO (1 mg/kg) for chemogenetic inhibition based on a previous report showing that a higher dose of CNO (5 mg/kg) could elicit non-specific response in the absence of DREADDs receptor expression [40]. Indeed, we did not observe any obvious effects in altering locomotion behaviors at the dose of 1 mg/kg following chemogenetic inhibition of dmPAG^Tac2^ neurons (**Figure S7A-D**), whereas in the elevated-plus maze test, except for an increase of time spent in the open arms (**Figure S7F**, see *Discussion*) after CNO administration, other measures for anxiety-like behaviors exhibited similar levels (**Figure S7E, G-H**). In summary, above results suggest that ablation or inhibition of dmPAG^Tac2^ neurons can reduce aggression behaviors in mice.

### Activation of dmPAG^Tac2^ neurons elicits fighting behaviors

Having confirmed the necessity of dmPAG^Tac2^ neurons in fighting behaviors from either ablation or inhibition of these cells, we then performed chemogenetic activation experiment to test the sufficiency of dmPAG^Tac2^ neurons in eliciting fighting behaviors. Neuron-specific Cre-dependent AAV encoding hM3Dq was injected into dmPAG of *Tac2*-Cre mice (**Figure 5A-B**, *Tac2*^hM3Dq^ mice). *Tac2*-Cre mice that received AAV-mCherry injections served as the control group (**Figure 5A-B**, *Tac2*^mCherry^). To confirm the efficacy and specificity of chemogenetic activation of dmPAG^Tac2^ neurons, CNO was administered (0.5 mg/kg) to examine c-Fos induction in Tac2^+^ cells (**Figure 5C**). The results revealed that c-Fos induction was specific to hM3Dq-expressing neurons but not to mCherry-expressing neurons (**Figure 5C-D**).

**Figure 5.**
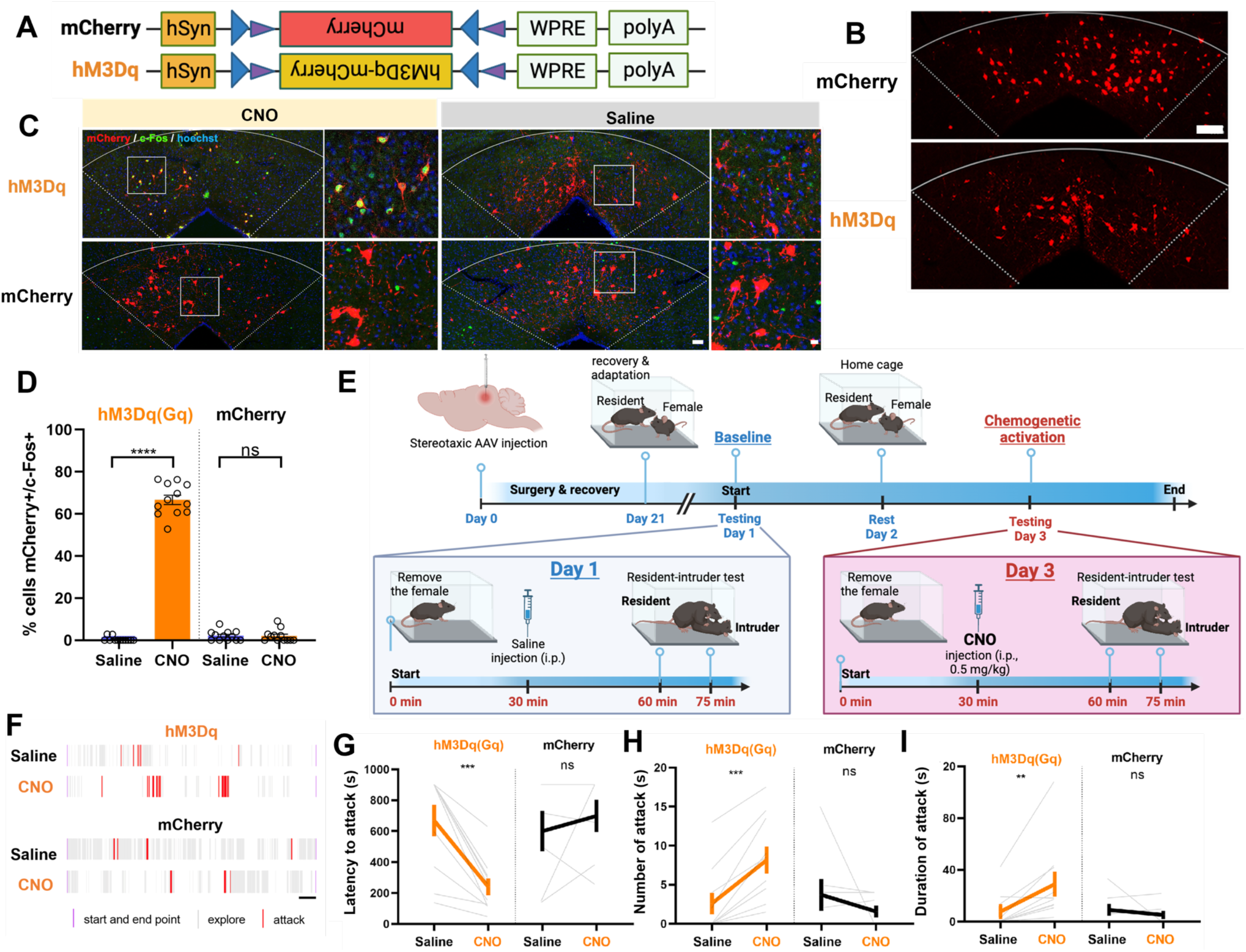
Chemogenetic activation of dmPAG^Tac2^ neurons elicits fighting behaviors. **(A)** A Cre-dependent hM3Dq vector or its control vector to be expressed in dmPAG^Tac2^ neurons. **(B)** Representative fluorescent images of hM3Dq-mCherry or mCherry (red) in dmPAG^Tac2^ neurons. Scale bar, 100 μm. **(C)** Activation of dmPAG^Tac2^ neurons in *Tac2*^hM3Dq^ and control *Tac2*^mCherry^ mice following intraperitoneal administration of CNO and normal saline, respectively (red: mCherry; green: c-Fos; blue: Hoechst). Scale bars, 50 μm (left) and 10 μm (enlarged right panel), respectively. **(D)** Number of mCherry^+^ cells in c-Fos^+^ neurons in *Tac2*-Cre mice receiving CNO injection (0.5 mg/kg, i.p.). N=3 mice, with 4 brain sections from each mouse. **(E)** Strategy for the RI test in *Tac2*-Cre mice receiving DREADDs vectors. **(F)** Raster plots showing mouse attack (red) or exploring behavior (gray) during the RI test. Time scale, 1 min. **(G-I)** Analysis of latency to attack **(G)**, number of attack **(H)**, and duration of attack **(I)**. N = 10 in *Tac2*^hM3Dq^ group and N = 7 in *Tac2*^mCherry^ group. Data represent mean ± S.E.M. Significance was calculated by means of paired *t*-test. **P < 0.01, ***P < 0.001, ****P < 0.0001.

Next, we examined the effects of chemogenetic activation of dmPAG^Tac2^ neurons on fighting behaviors. First, we established the baseline level of aggression by administering *Tac2*-Cre mice with normal saline and subjecting them to the RI test after waiting for 30 min. Forty-eight hours later, these mice were intraperitoneally injected with CNO (0.5 mg/kg) and subjected to the RI again after waiting for 30 min (**Figure 5E**). Upon saline administration, *Tac2*^mCherry^ mice had no changes in fighting behaviors, similar to the results of *Tac2*^mCherry^ mice upon chemogenetic inhibition (**Figure 4L-N**). However, following CNO administration, *Tac2*^hM3Dq^ mice exhibited a significantly elevated levels of aggression towards the intruder (**Figure 5F-I**). There was a 3-fold increase in percentage of mice exhibiting fighting behaviors following activation of dmPAG^Tac2^ neurons (**Figure S6D**). In addition, we tested the possible effects of CNO administration on other behavioral performances and failed to observe any obvious effects of chemogenetic activation of dmPAG^Tac2^ neuron in altering locomotion (**Figure S8A-D**) and anxiety-like behaviors (**Figure S8E-H**). In summary, activation of dmPAG^Tac2^ neurons elicits fighting behaviors and increases aggression in male mice without producing obvious effects in the other behaviors tested.

### Fighting behaviors alters transcriptional profiles in dmPAG^Tac2^ neurons

The above data demonstrate that the dmPAG^Tac2^ neurons not only are involved, but also play a causal role in fighting behaviors measured by the RI. Next, we aimed to reveal the molecular programs in dmPAG^Tac2^ neurons that are altered by fighting behaviors. To this end, we performed TRAP by injecting Cre-dependent AAV vector with eGFP-tagged ribosome protein L10α (**Figure 6A-B**) in the dmPAG of *Tac2*-cre mice. These mice were subjected to the RI or were maintained as home-cage controls (**Figure 6C**, see *Methods*). Sixty minutes after the neurons were purified and RNA-sequencing (RNA-seq) were performed. By comparing gene expression between the two groups of immunoprecipitated (IP) samples (RI vs Control), we identified 58 differentially expressed genes (DEGs) induced by fighting, including 34 upregulated DEGs and 23 downregulated DEGs (**Figure 6D**). Kyoto Encyclopedia of Genes and Genomes (KEGG) analysis showed that, the pathways impacted by fighting behaviors mainly involved inflammation, immunity and infection-related pathways (**Figure 6E-F**). Of note, the estrogen signaling pathway, which has been previously shown in controlling inter-male aggression behaviors in the VMH [11], was identified by our analysis, consistent with the previous notion that estrogen can regulate aggression behaviors [21, 41, 42]To identify novel targets related to fighting, we performed enrichment analysis [43] by calculating fold-enrichment (RI-IP/Ctr-IP) and aligned the top candidate genes (**Figure 6G**). Among the most significantly enriched genes, several well-established immediate-early genes, including *Arc*, *Egr1*, *Fgf2*, and *Fosb*, were upregulated, suggesting that, indeed, the dmPAG^Tac2^ neurons were activated by fighting behaviors and that the TRAP-seq approach was accurate in capturing these molecular events. In downregulated genes, we noticed that *Tph2*, a gene encoding tryptophan hydroxylase 2, a rate-limiting enzyme that is primarily expressed in the serotonergic neurons of the brain, consistent with that previously established evidence showing that the serotonergic system is involved in aggression at both ligand and receptor levels [44–47].

**Figure 6.**
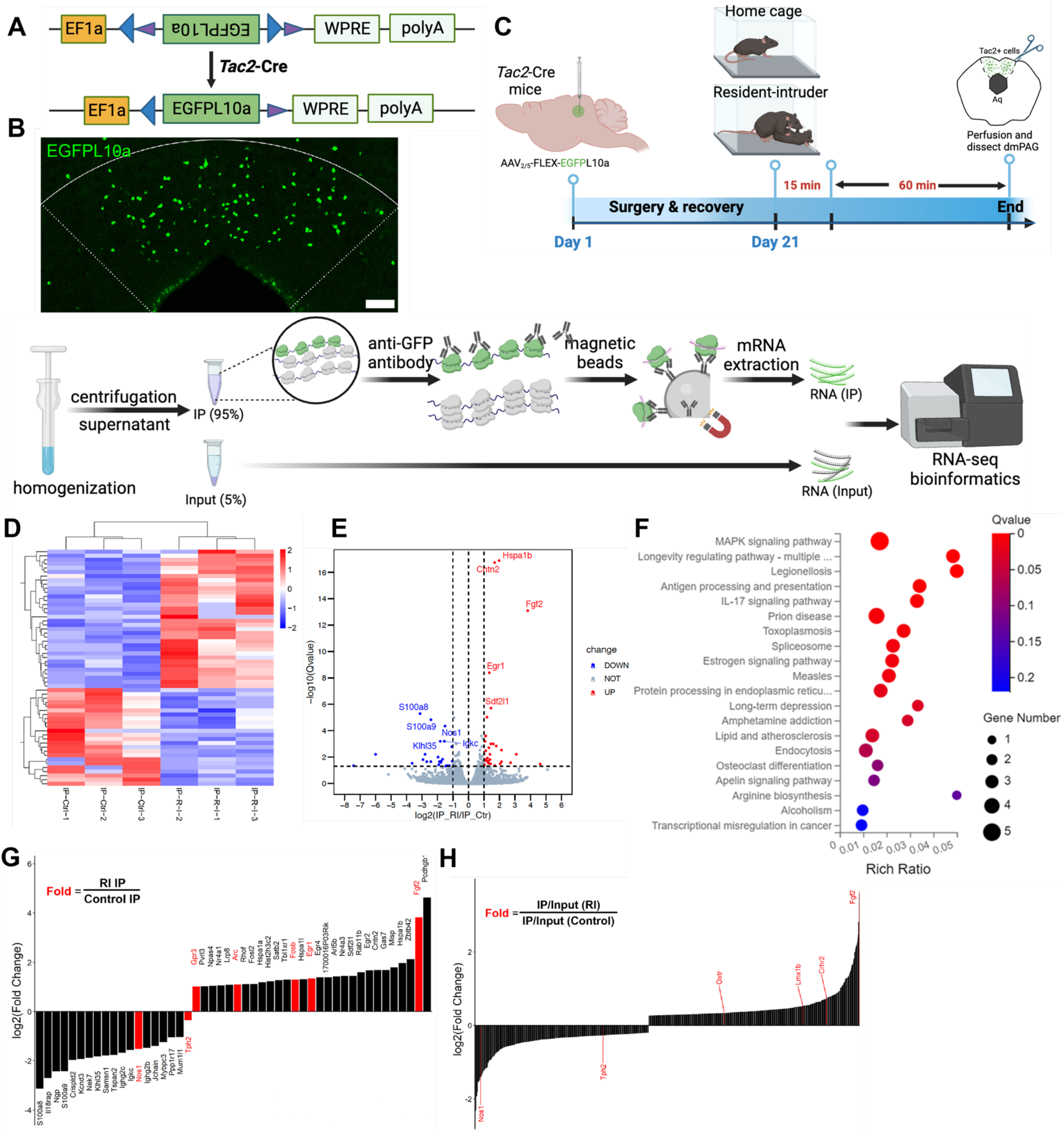
TRAP-seq reveals transcriptional alterations following fighting behaviors. **(A)** A Cre-dependent TRAP (AAV-FLEX-EGFPL10a) vector to be expressed in *Tac2*-Cre mice. **(B)** Representative fluorescent images of EGFPL10a protein (green) in dmPAG. Scale bar, 100 μm. **(C)** Strategy for TRAP-seq in *Tac2*-Cre mice injected with AAV-FLEX-EGFPL10a in dmPAG and subjected to the RI assay or maintained at home cage (control). **(D)** Heatmap showing DEGs of the IP samples. |Log2FC|>1, adjusted q value < 0.05. **(E)** Volcano plot showing DEGs of the IP samples. **(F)** KEGG enrichment analysis showing top 20 most enriched pathways affected by RI when compared to the control. **(G)** Fold-change analysis showing log2(FC) value of RI-IP to Control-IP samples. Genes labeled in red are immediate-early genes (*Gpr3*, *Arc*, *Fosb*, *Egr1*, *Fgf2*) or genes related to the 5-HT system. **(H)** A secondary analysis of enrichment score, based on normalizing IR/Input (RI) to IR/Input (Control). Note that *Nos1*, *Tph2*, *Oxtr*, *Lmx1b*, *Crhr2*, and *Fgf2* (labeled in red on the left) were listed among the most enriched genes that were related to the 5-HT system.

### Pharmacological manipulation of dmPAG^Tac2^ neurons in fighting behaviors

To validate the potential molecular pathways in dmPAG^Tac2^ neuron altered by fighting, we targeted the 5-HT system as identified by TRAP-seq. We hypothesize that increasing 5-HT levels in the extracellular synaptic cleft by intervening 5-HT reuptake, as previously demonstrated [48, 49], could block the behavioral effects induced by chemogenetic activation of dmPAG^Tac2^ neurons. To this end, we intracranially delivered fluoxetine, a widely used selected 5-HT reuptake inhibitor, into the dmPAG prior to chemogenetic activation (see *Methods*, **Figure 7A**). As expected, in *Tac2*^hM3Dq^ mice, CNO administration effectively elicited fighting while intra-dmPAG delivery of fluoxetine prevented the chemogenetic activation-elicited fighting behaviors, as showed by a significantly increased latency to attack (**Figure 7C**), decreased number (**Figure 7D**) and duration of attacks (**Figure 7E**), suggesting that targeting 5-HT reuptake in dmPAG neurons could prevent the fighting behaviors elicited by activation of dmPAG^Tac2^ neurons. The above data suggest potential involvement of 5-HT system in dmPAG^Tac2^ neurons-mediated fighting behaviors. We also analyzed the TRAP-seq data mainly focused on 5-HT system-associated genes, including genes that encode 5-HT receptors (**Figure S9)**. The analysis shows RI indeed upregulated gene expression of 5-HT receptors (**Figure S9**).

**Figure 7.**
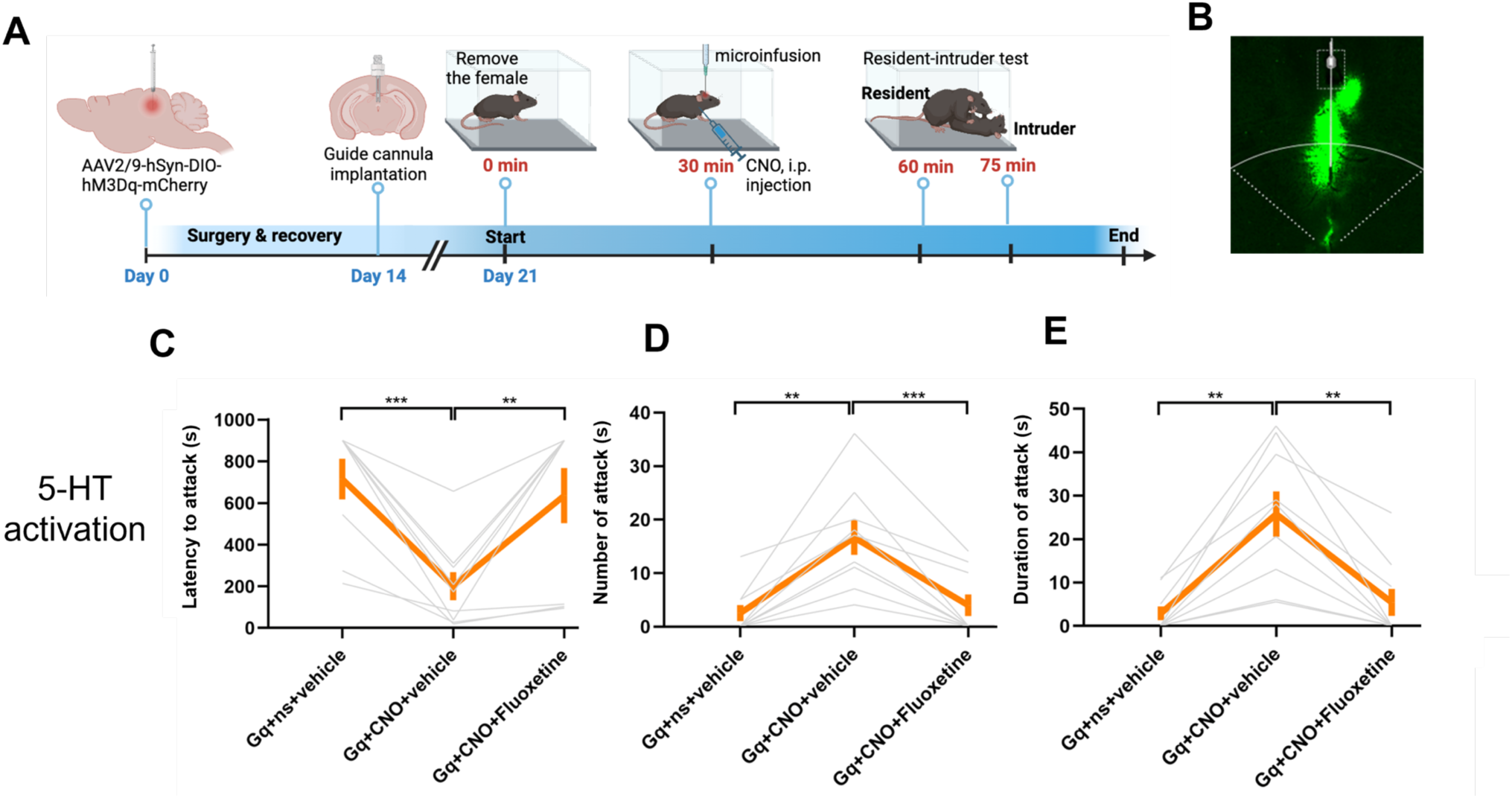
Pharmacological validation of molecular targets identified by TRAP-seq. **(A)** Strategy for activation of the 5-HT system with fluoxetine microinfusion to suppress fighting behaviors induced by chemogenetic activation of dmPAG^Tac2^ neurons. **(B)** Representative fluorescent image showing cannula position targeting dmPAG region in *Tac2*-Cre mice. **(C-E)** Analysis of latency to attack **(C)** number of attack **(D)** and duration of attack **(E)** in *Tac2*^hM3dq^ mice injected with fluoxetine/vehicle and CNO/saline at dmPAG. N = 9 mice/group. Data represent mean ± SEM. Significance was calculated by means of paired *t*-test. * P < 0.05, ** P < 0.01, *** P < 0.001, **** P < 0.0001.

## Discussion

### dmPAG^Tac2^ neurons exhibit distinct expression pattern

Tac2^+^ neurons showed a unique distribution pattern in the mouse brain. At basal level, *Tac2* genes are expressed in a wide variety of brain areas associated with mood and social behaviors, including the bed nuclei of the stria terminalis (BNST), medial habenular (MHb), central amygdala (CeA) [10]. In pathological states such as prolonged social isolation, Tac2 can be strongly induced to act on the brain in a dissociable and region-specific manner [10]. The induction pattern of Tac2 peptide or distributed expression of Tac2^+^ neurons suggest that the regulatory roles of the Tac2/NkB signaling in multiple brain functions depend on its anatomical location and neural connection. For instance, *Tac2*-expressing neurons in the dorsomedial shell of the nucleus accumbens have been previously characterized as a subtype of D1 medium spiny neurons that play a negative regulatory role in cocaine addiction [50]. In CeA, *Tac2*-expressing neurons mainly modulate consolidation of fear memory [9, 51]. A recent study showed that *Tac2*-expressing neurons in the BNST and the CeA regulate behavioral states, i.e., emotional valence, by sending inhibitory inputs to the LHA neurons expressing melanin-concentrating hormone [52]. These studies suggest that *Tac2*-expressing neurons are uniquely positioned in specific brain regions to modulate a variety of behavioral functions.

However, the distribution pattern of Tac2 in the midbrain and the specific functions of Tac2 neurons in corresponding brain regions are not fully understood. In this study, we focus on *Tac2*-expressing neurons located in the midbrain and revealed that Tac2^+^ neurons were specifically distributed in the PAG, with a higher expression level in the dorsomedial part (dmPAG).The PAG has been traditionally viewed as an integrating center for defensive responses, including panic, fight-flight and freezing behaviors [1, 7]. Early studies using c-Fos mapping revealed that there was an anatomic segregation of defensive behaviors in the dorsal columns of the PAG [53] and that the dmPAG is served as a relay station for acquiring aversive information to the other brain regions [54]. However, there is a lack of study on whether there was a molecularly distinct cell type that might mediate this effect. Here, we uncovered that *Tac2*-expressing neurons in the dmPAG (dmPAG^Tac2^ neurons) are a key cell type that plays a critical role in intermale aggression.

### dmPAG^Tac2^ neurons are recruited during intermale aggression

Having confirming that *Tac2*-expressing neurons in the PAG are mainly located in the dmPAG, we hypothesize that dmPAG^Tac2^ neurons are involved in aggression behaviors. Using a RI paradigm, we found that fighting behaviors specifically activate dmPAG^Tac2^ neurons, which are shown by induction of c-Fos and time-locked Ca^2+^ transient as showed by our fiber photometry experiments (**Figure 3**), suggesting that dmPAG^Tac2^ neurons are recruited during aggression.

Previous studies have demonstrated a sex-specific effect of Tac2^+^ neurons in the CeA [8, 21, 51]. Whether this effect pertains in the dmPAG^Tac2^ neurons, however, is not clear. While we only assessed the role of dmPAG^Tac2^ neurons in regulation of intermale aggression, future studies should also investigate whether these neurons play a role in other types of aggression such as maternal aggression and predatory aggression. In fact, the PAG is also known as a center in regulating predatory aggression [55], in which the lPAG mainly recruited for sensory processing [56] and the vlPAG glutamatergic neurons recruited for conditioned freezing responses [57]. However, whether dmPAG^Tac2^ neurons are directly or indirectly involved in predatory aggression is currently unknown and will be an interesting research topic for future study.

### Bidirectional manipulation of dmPAG^Tac2^ neuronal activity alters fighting behaviors

Our targeted inhibition and genetic ablation of dmPAG^Tac2^ neuronal activity yielded qualitatively similar results in RI (**Figure 4**), showed by prolonged the latency to attack and significantly reduced numbers and duration of attacks, suggesting suppressed fighting behaviors. Thus, dmPAG^Tac2^ neurons are necessary to mediate fighting behaviors. On the other hand, targeted chemogenetic activation of dmPAG^Tac2^ neuronal activity, however, not only shortened the latency to attack, but also significantly increased numbers and duration of attacks (**Figure 5**), demonstrating that activation of dmPAG^Tac2^ neurons is sufficient to induce fighting behaviors. This striking evidence shows that dmPAG^Tac2^ neurons play a critical role in fighting behaviors in male mice. In chemogenetic manipulation, we chose a before-after study design based on several previous studies [11, 22, 25] to maximally measure the effect of interventions on the same subjects, providing an evaluation of variables at baseline and after interventions [58].

### Role of 5-HT systems in dmPAG^Tac2^ neurons during aggression

The relationship between the 5-HT system and aggression has been a significant area of study. For instance, male mice with a mutation in the monoamine oxidase A gene show enhanced aggression accompanied by altered brain 5-HT levels [59, 60]. Similarly, mice lacking the 5-HT_1B_ receptor exhibited intense fighting behaviors towards an intruder [61], and mice with global nNOS knockout exhibited high levels of aggression, associated with impaired 5-HT turnover and deficient 5-HT receptor function [62]. In our study, while we observed some evidence suggesting a potential involvement of the 5-HT system in dmPAG^Tac2^ neurons-mediated fighting behaviors, the results are preliminary and indicate a possible role of the 5-HT system in this molecularly distinct neuronal subtype.

Using TRAP-seq, we focused on the translational alterations in *Tac2*-expressing neurons in the dmPAG following RI exposure in mice that were screened for their aggressiveness. Instead of directly studying Tac2/NkB signaling or Tac2 peptide, our investigation targeted the 5-HT system. We found that inhibition of 5-HT reuptake with fluoxetine could suppress fighting behaviors elicited by activating dmPAG^Tac2^ neurons. While these results provide some validation for the targets revealed by TRAP-seq, they are preliminary and should be interpreted cautiously. These findings align with earlier reports implicating that the 5-HT system in intermale aggression [59–63], but further research is necessary to fully understand these pathways.

### Limitations

In clinical assessment, aggression behaviors are predominantly categorized into two major subtypes, impulsive aggression or premeditated aggression [64], however, in animal studies, aggressive and defensive behaviors are characterized into more broad categories, including play fighting, offensive aggression, defensive aggression, predatory aggression [23] [65]. One of the limitations of our study is, we mainly assess the role of dmPAG^Tac2^ neurons in intermale aggression, but not in other subtypes of aggressive behaviors such as female aggression, maternal aggression, and predatory aggression, as discussed above. Recent studies have demonstrated a sex-dependent role of Tac2^+^ neurons in the regulation of fear memory [51], suggesting the importance of consideration of sexual dimorphism.

### Potential treatment for humans with exaggerated aggression

Uncontrolled aggression in humans poses a significant threat to society, leading to immediate physical harm can causing chronic stress that can negatively impact mental health [66]. In extreme cases, human aggression can escalate into incidents such as homicide or mass shooting [67]. Therefore, finding effective ways to control aggression behaviors is crucial. In this study, we identify a molecularly distinct neuronal cell type in the midbrain that plays a role in controlling intermale aggression. While our findings suggest potentially druggable targets to modulate exaggerated aggression, it is important to note that our results primarily identifies a druggable cell type rather than specific genetic targets. Additionally, the resident intruder paradigm, used as a model for aggression, may not fully capture the complexity of human aggression. This limitation should be considered when interpreting the potential translational impact of our findings.

## Acknowledgement

This work was funded by National Natural Science Foundation of China (No. 82173197 and 81972362) to X.Q.C. The authors thank Prof. Minmin Luo for kindly providing *Tac2*-Cre knock-in mouse line, Prof. Jian-Zhi Wang for kindly providing Ai9/tdTomato mice, and Prof. Yunyun Han for kindly providing suggestions on Ca^2+^ imaging. The authors gratefully thank the members from the X.Q. Chen Lab help with this project during the first-wave of COVID-19 in Wuhan.

## Author contributions

C.Y.L. researched, analyzed, interpretated data, and helped write the manuscript. C.M., J.W., Y.G., P.G., S.Y., P.Z., and B.T. researched data and helped analyze data. W.C. conceived and designed the study, interpretated data and wrote the manuscript; X.Q.C. secured the funding, designed the study, interpretated data, supervised all work and helped write the manuscript.

## Competing interests

The authors declare no competing interests.

## Abbreviations

5-HT: 5-hydroxytryptamine
7-NI: 7-Nitroindazole
AAV: adeno-associated virus
ACC: anterior cingulate cortex
AHN: anterior hypothalamic nucleus
ARH: arcuate hypothalamic nucleus
BLA: basolateral amygdalar nucleus
BNST: bed nuclei of the stria terminalis
CeA: central amygdalar nucleus
CNO: clozapine N-oxide
CUN: cuneiform nucleus
DEPC: diethypyrocarbonate
dIPAG: dorsolateral periaqueductal gray
DMH: dorsomedial nucleus of the hypothalamus
dmPAG: dorsomedial periaqueductal gray
DMSO: dimethyl sulfoxide
DR: dorsal nucleus raphe
DTA: diphtheria toxin
EDTA: ethylene diamine tetraacetic acid
EGFP: enhanced green fluorescent protein
EPM: elevated plus maze
GABA: γ-aminobutyric acid
HPF: hippocampal formation
LHA: lateral hypothalamic area
LPAG: lateral periaqueductal gray
MeA: medial amygdalar nucleus
MHb: medial habenula
MM: medial mammillary nucleus
MPN: medial preoptic nucleus
MPO: medial preoptic area
MRN: midbrain reticular nucleus
NAc: nucleus accumbens
NK: neurokinin
nNOS: neuronsal nitric oxide synthase
OB: olfactory bulb
OCT: optimal cutting temperature compound
OFT: open field test
PAG: periaqueductal gray
PBS: phosphate buffer saline
PVT: paraventricular nucleus of the thalamus
SSRI: selective serotonin reuptake inhibitor
Tac: tachykinin
TPH2: tryptophan hydroxylase2
TRAP: translating ribosome affinity purification
VIPAG: ventrolateral periaqueductal gray
VMH: ventromedial hypothalamic nucleus

## Supplementary Figure Legends

**Figure S1. PAG *Tac2*-expressing neurons are mainly located in the dmPAG (related to Figure 1).**

**(A)** Representative fluorescent images showing absence of tdTomato-labeled Tac2^+^ cells in control mice (*Tac2*-Cre, Ai9, or WT C57BL/6 mice). Scale bars, 500 μm. **(B)** *Tac2* mRNA expression pattern in ABA ISH database (left), with an enlarged view (right) from the left panel. **(C)** Distribution pattern of Tac2^+^ cells in PAG subregions examined fluorescence ISH. Scale bars, 50 μm. **(D)** Pie chart showing numbers of *Tac2*^tdTomato^ cells in PAG subregions. N = 3 mice. Data were expressed as mean ± S.E.M.

**Figure S2. Mapping of c-Fos expression in WT mice subjected to the RI assay (related to Figure 2)**

**(A)** Brain areas with high-level c-Fos expression in WT mice subjected to the RI assay. Scale bar, 1,000 μm. **(B)** Representative fluorescent image showing multiple hypothalamic subnuclei were activated by the RI assay, including the PVT, SBPV, PVZ, AHN, LHA, BLA, DMH, VMH, MRN, and PAG. Scale bar, 100 μm. **(C)** Quantification of number of Fos+ cells in selected brain areas in WT mice subjected to the RI assay. Data are presented as total number of Fos+ cells in five brain sections separated by 200 μm (N = 3 mice per condition). Data were expressed as mean ± S.E.M. *Abbreviations*: PVT, Paraventricular nucleus of the thalamus; SBPV, Subparaventricular zone hypothalamus; PVZ, Periventricular zone; AHN, Anterior hypothalamic nucleus; LHA, Lateral hypothalamic area; BLA, Basolateral amygdala nucleus, DMH, Dorsomedial nucleus of the hypothalamus; VMH, Ventromedial hypothalamic nucleus; MRN, Midbrain reticular nucleus; PAG, Periaqueductal gray.

**Figure S3. Using a Cre-out strategy to test the role of dmPAG^Tac2^ negative neurons in aggression (related to Figure 3)**

**(A)** Schematic showing a Cre-out strategy. **(B-D)** Ca^2+^ activity time-locked to each attack episode. The panels showing **(B)** Distribution of attack episode, **(C)** Heatmap of GCaMP6m, and **(D)** ΔF/F%. **(E)** Analysis of peak ΔF/F before (Pre) and after (Post) each attack episode. Data represent mean ± S.E.M. Significance was calculated by means of paired *t*-test. \**P* < 0.05. **(F)** Analysis of peak ΔF/F in dmPAG^Tac2^ neurons (N = 10 trails, in Figure 3H) versus dmPAG^Tac2^ negative neurons (N = 46 trails) during each attack episode. Data represent mean ± S.E.M. unpaired *t*-test, \*\*\*\**P* < 0.0001.

**Figure S4. Tac2-expressing neurons in dmPAG respond to aggression (related to Figure 3)**

**(A-F)** Ca^2+^ activity time-locked to mice encounter **(A-B)**, male-female encounter **(C-D)**, and male-female interaction **(E-F)**. **(A)** Heatmap of GCaMP6m ΔF/F signals (top) and ΔF/F% (bottom). **(B)** Analysis of peak ΔF/F before (Pre) and after (Post) each male-male encounter episode. The other panels present the same manner but with different behavioral events, including male-female encounter **(C-D)** and male-female interaction **(E-F). (G-H)** Ca^2+^ activity time-locked to motor activity in the rotarod test. **(G)** Heatmaps of GCaMP6m ΔF/F signals (top) and ΔF/F% (bottom). **(H)** Analysis of peak ΔF/F before (Pre) and after (Post) the start of rotarod test. **(I-P)** Ca^2+^ activity time-locked to object exploration **(I-J)**, fox odor sniffing **(K-L)**, saline odor sniffing **(M-N)**, and pain stimulation **(O-P)**. **(I)** Distribution of object exploration (top), heatmaps of GCaMP6m ΔF/F signals (middle) and ΔF/F% (bottom). **(J)** Analysis of peak ΔF/F before (Pre) and after (Post) each object exploration episode. The other panels present the same manner but with different behavioral events, including fox odor sniffing **(K-L)**, saline odor sniffing **(M-N)**, and pain stimulation **(O-P)**. Data represent mean ± S.E.M. Significance was calculated by means of paired *t*-test.

**Figure S5. Behavioral phenotypes (locomotion, anxiety, social behaviors) in *Tac2*-Cre mice (related to Figure 4).**

**(A)** Schematic of the open-field test. **(B-E)** Locomotor performance of WT and *Tac2*-Cre mice on **(B)** total distance, **(C)** average speed, **(D)** time spent in central and peripheral zones, and **(E)** entries into the central and peripheral zones. **(F)** Schematic of the elevated-plus maze test. **(G-J)** Anxiety-like behaviors of WT and *Tac2*-Cre mice on **(G)** numbers of entries in closed arm, **(H)** time spent in closed arm, **(I)** numbers of entries in open arm, **(J)** time spent in open arm. **(K)** Schematic of the social interaction test (sociability session). **(L-M)** Social preference of WT and *Tac2*-Cre mice on **(L)** interaction time and **(M)** social preference index. **(N)** Schematic of the social interaction test (social novelty session). **(O-P)** Social preference of WT and *Tac2*-Cre mice on **(O)** interaction time and **(P)** social preference index. **(Q)** Schematic of the RI assay. **(R-T)** Behavioral performance of WT and *Tac2*-Cre mice on **(R)** latency to attacks, **(S)** number of attacks and **(T)** duration of attacks. Data represent mean ± S.E.M. N = 24 in WT mice and N = 25 in *Tac2*-Cre mice. Unpaired t-test, * *P* < 0.05, *** *P* < 0.001, **** *P* < 0.0001.

**Figure S6. Percentage of *Tac2*-Cre mice exhibit attack behaviors (related to Figure 4-5).**

**(A)** Percent of mice exhibiting attack before and after genetic ablation with DTA or mCherry (control). (**B**) Number (left) and duration (right) of investigation before and after ablation of dmPAG^Tac2^ neurons with DTA in *Tac2*-cre mice. N = 7 mice. Data represent mean ± S.E.M. Significance was calculated by means of paired *t*-test. **(C)** Percent of mice exhibiting attack before and after receiving hM4Di or mCherry (control). N = 10 in *Tac2*^hM3Dq^ and N = 7 in *Tac2*^mCherry^ and N = 7 in *Tac2*^mCherry^ group. Data represent mean ± S.E.M. Significance was calculated by One-way ANOVA and Tukey’s multiple comparisons test,

**Figure S7. Behavioral phenotypes (locomotion and anxiety behaviors) following chemogenetic inhibition (related to Figure 4).**

**(A-D)** Performance of the open-field test in *Tac2*^hM4Di^ mice receiving saline or CNO (1 mg/kg, *i.p.*). Analysis showing results of total distance **(A)**, average speed **(B)**, time spent in the central or peripheral zones **(C)**, and entries into the central or peripheral zones **(D)**. **(E-H)** Performance of the elevated-plus maze in *Tac2*^hM4Di^ mice receiving saline or CNO (1 mg/kg, *i.p.*). Analysis showing results of entries in the open arm **(E)**, total time spent in the open arm **(F)**, entries in the closed arm **(G)**, and total time spent in the closed arm **(H)**. N = 10 mice. Data represent mean ± S.E.M. Significance was calculated by means of paired *t*-test, \**P* < 0.05.

**Figure S8. Behavioral phenotypes (locomotion and anxiety behaviors) following chemogenetic activation (related to Figure 5).**

**(A-D)** Performance of the open-field test of *Tac2*^hM3Dq^ mice receiving saline or CNO (0.5 mg/kg, *i.p.*). Analysis showing results of total distance **(A)**, average speed **(B)**, total time in the central or peripheral zones **(C)**, and entries in the central or peripheral zones **(D)**. **(E-H)** Performance of elevated-plus maze of *Tac2*^hM3Dq^ mice receiving saline or CNO (0.5 mg/kg, *i.p.*). Analysis showing results of entries in the open arm **(E)**, total time spent in the open arm **(F)**, entries in the closed arm **(G)**, and total time spent in the closed arm **(H)**. N = 8 mice. Data represent mean ± S.E.M. Significance was calculated by means of paired *t*-test.

**Figure S9: Pharmacological validation of molecular targets identified by TRAP-seq (related to Figure 7)**

**(A)**. Heatmap showing counts of *Tph2* from TRAP-seq raw results. **(B)** Heatmap showing Log_2_FC value of *Tph2* transcript counts in individual sample pairs. **(C)** Counts of 5-HT system-related genes detected in the IP samples from RI mice and home-cage controls. **(D)** Log_2_FC value in the counts of 5-HT system-related genes in the IP samples from RI mice normalized to that of the home-cage control mice.

## Notes

### Competing Interest Statement

The authors have declared no competing interest.

### Summary of Updates

This manuscript has been peer-reviewed several times, and we have received very helpful comments. We have clarified and revised the manuscript. This is the version #3 of formal revision. We hope to provide an early access to the general community.

