## Supplementary figures and images for "A molecularly distinct cell type in the midbrain regulates intermale aggression behaviors in mice"

### Supplemental figures

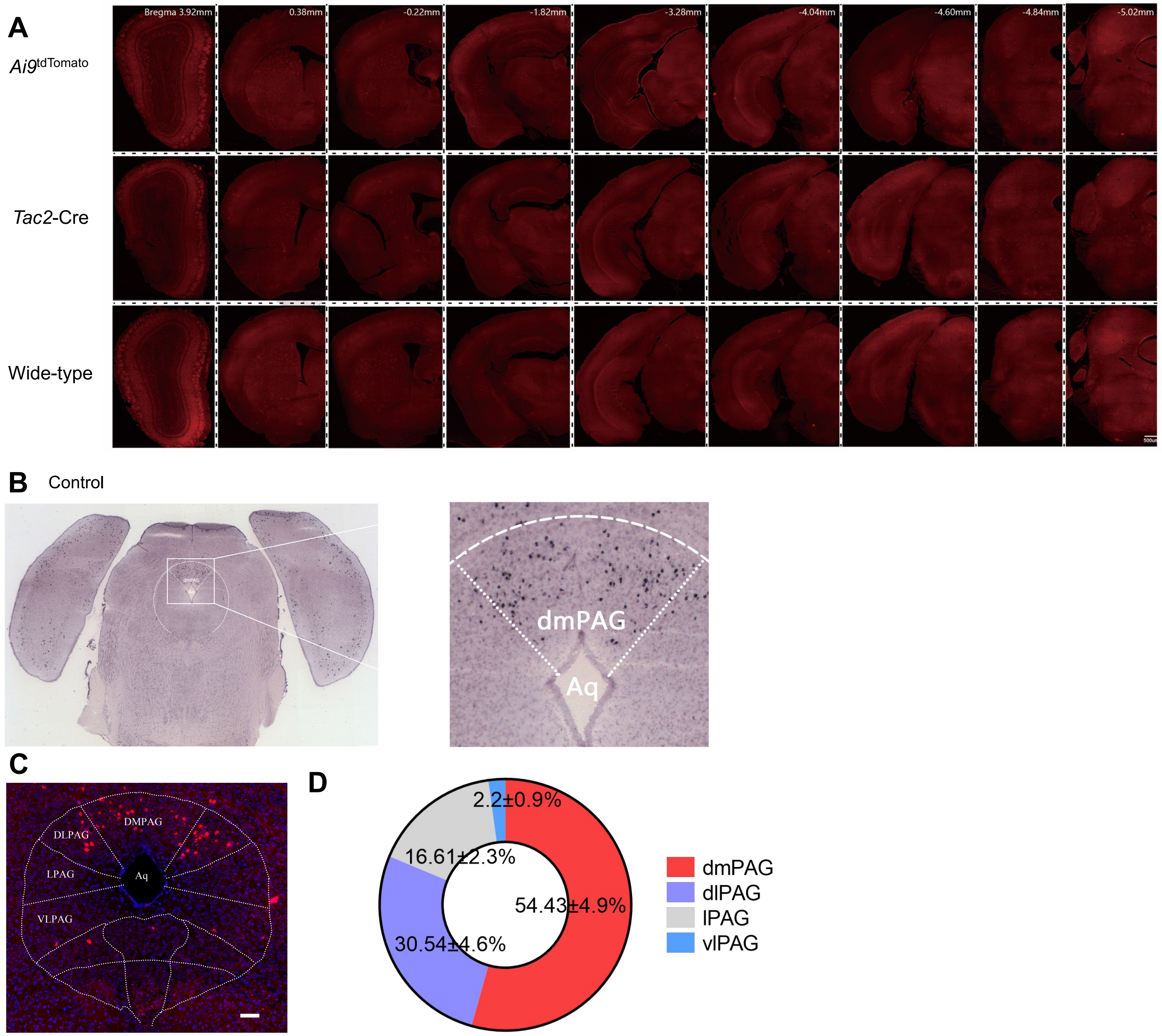

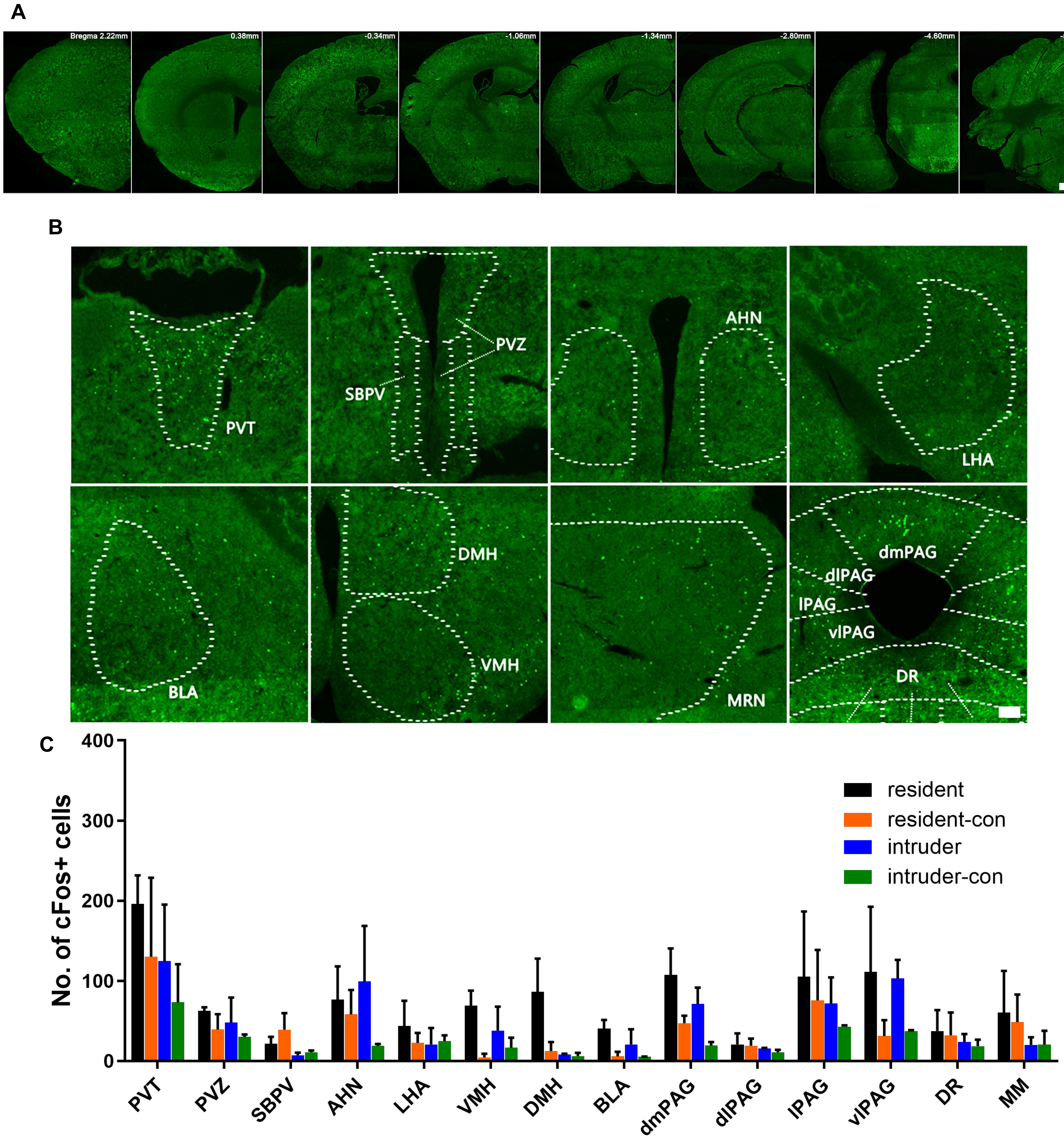

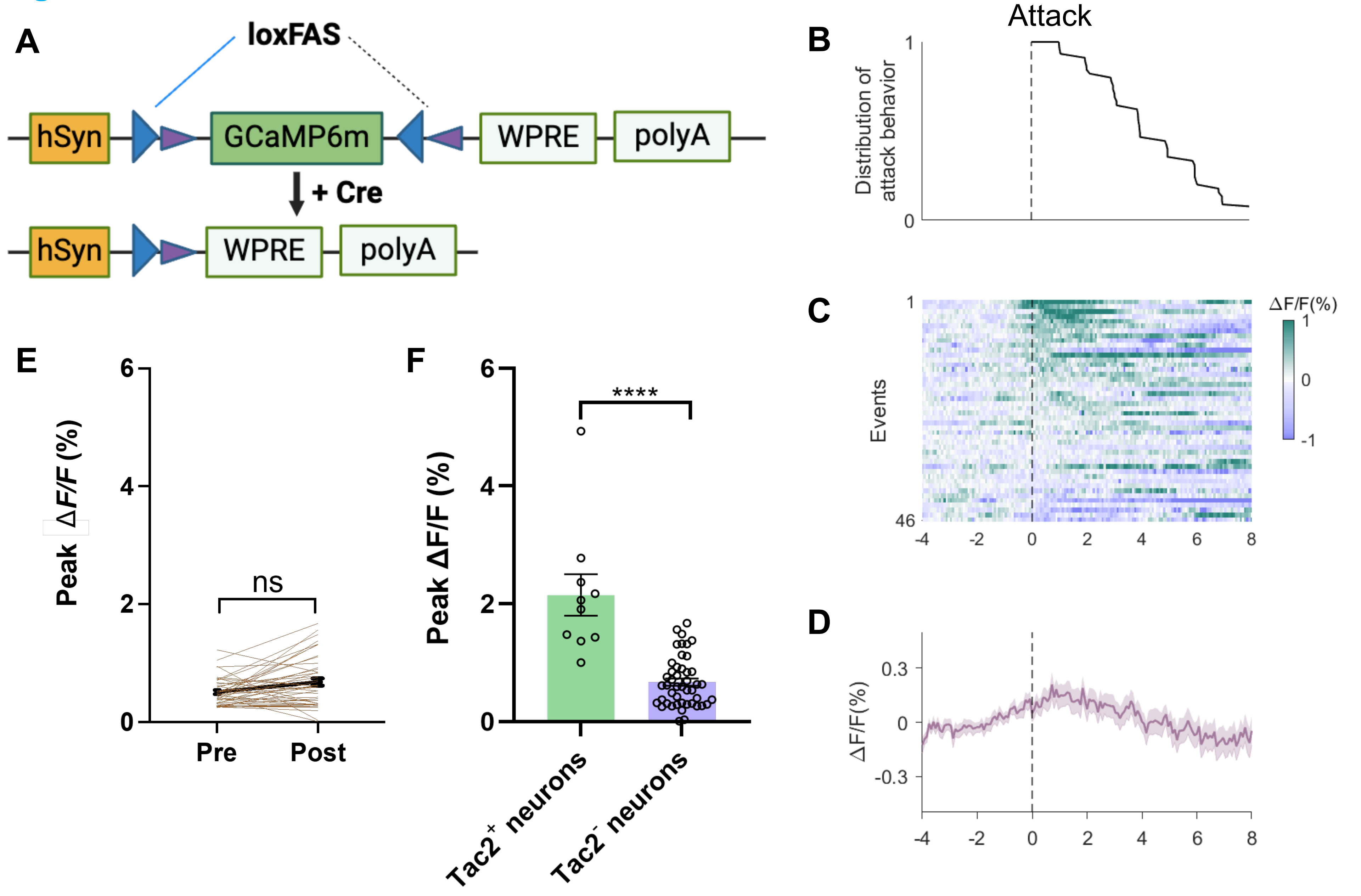

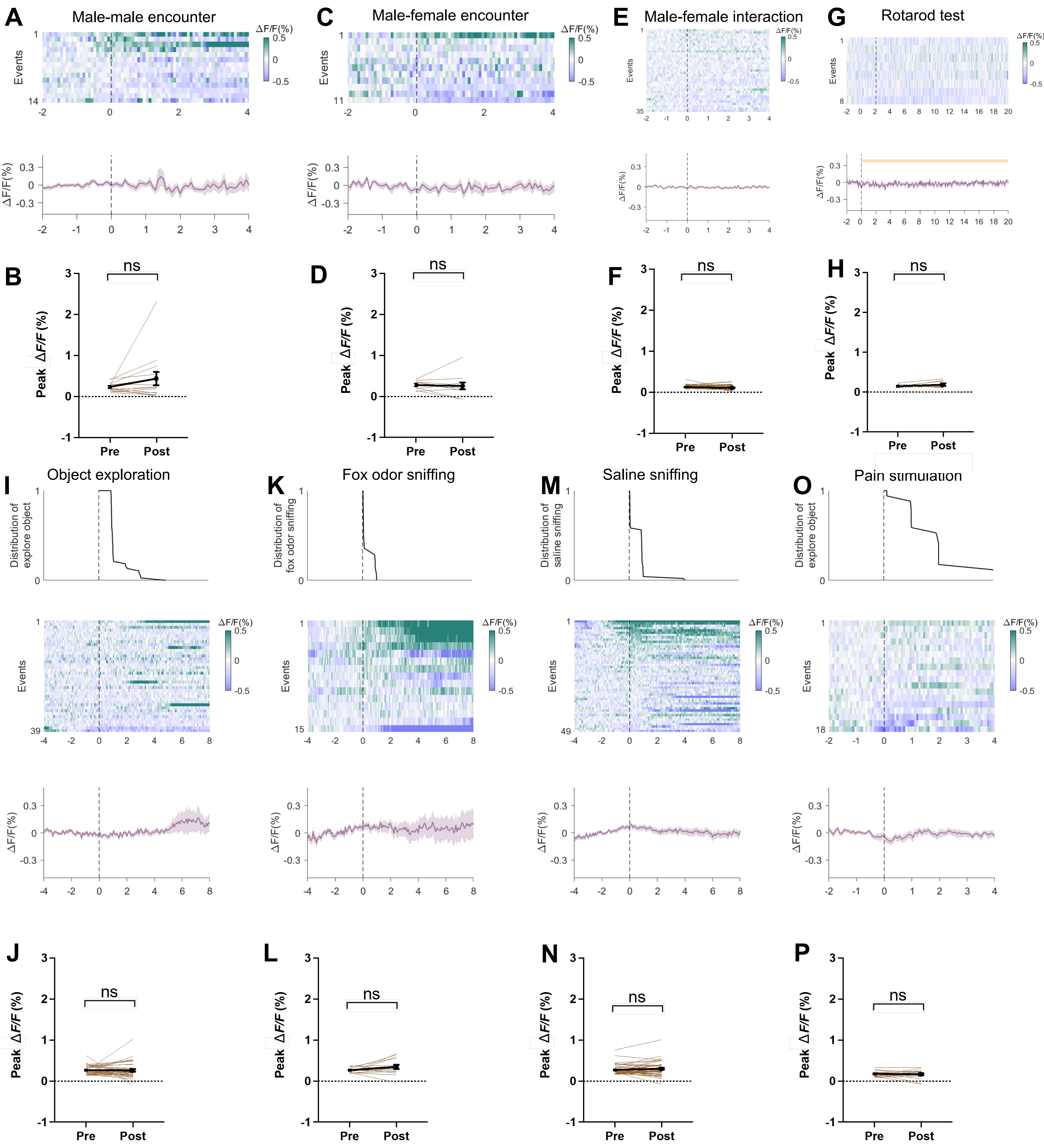

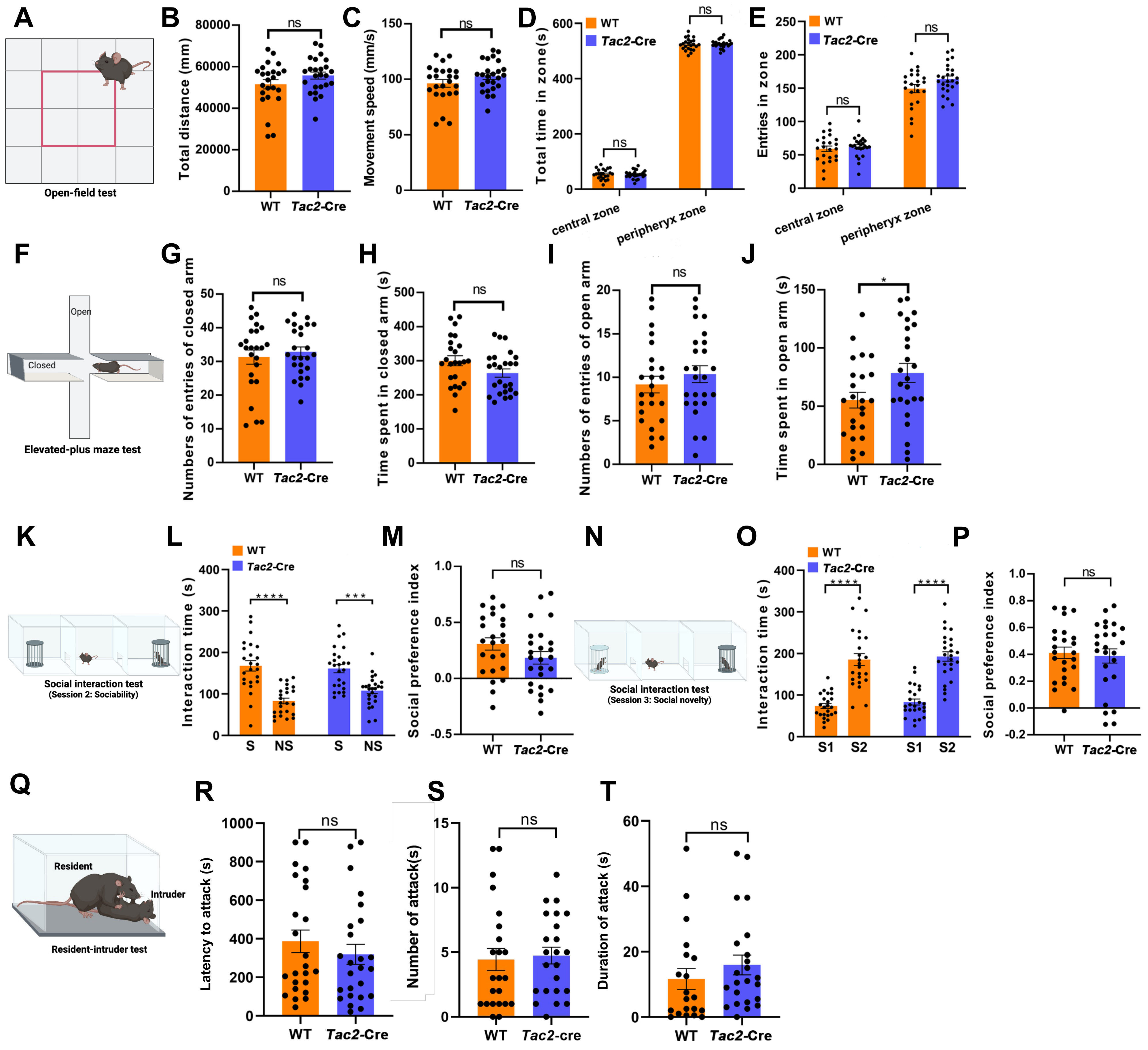

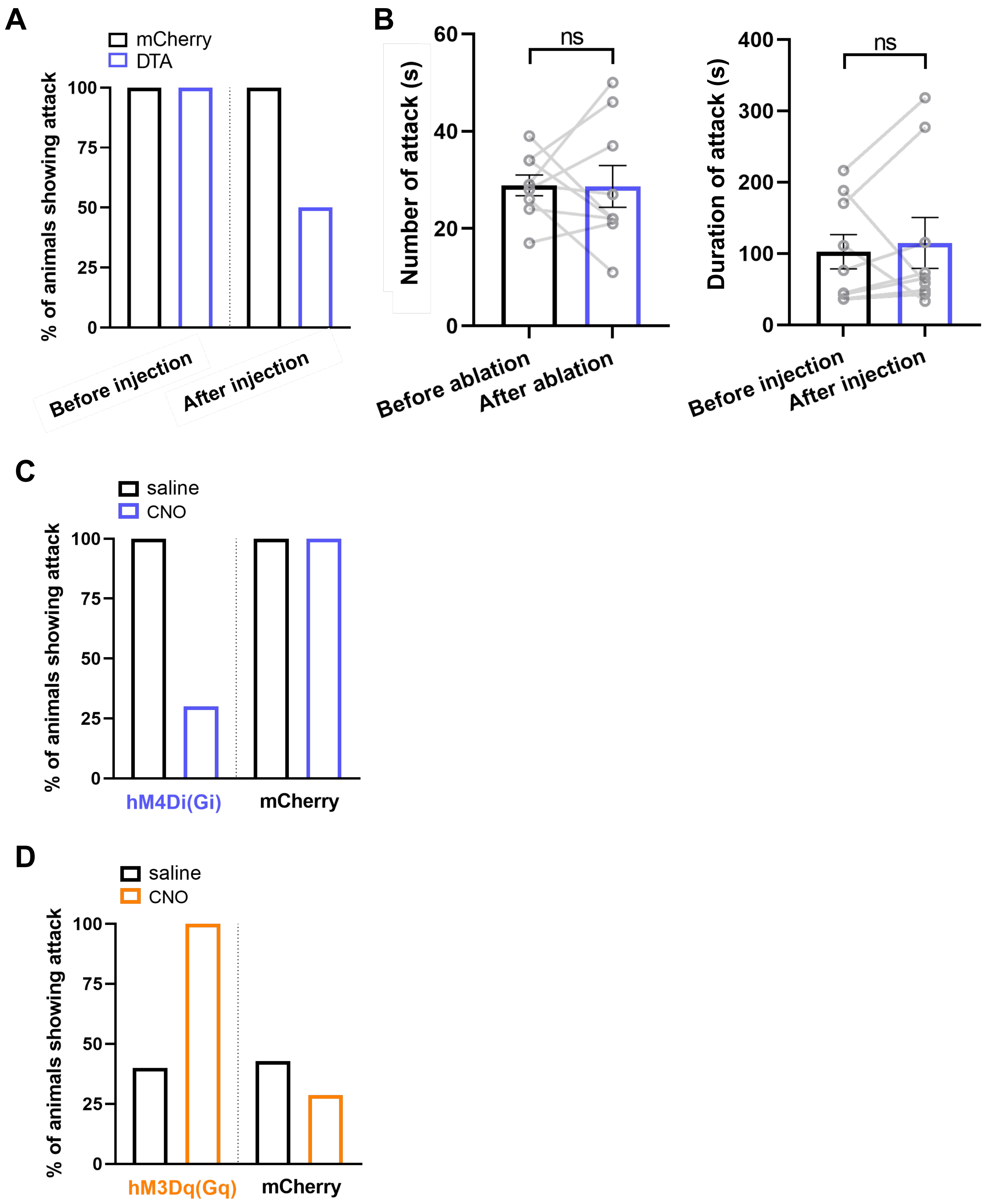

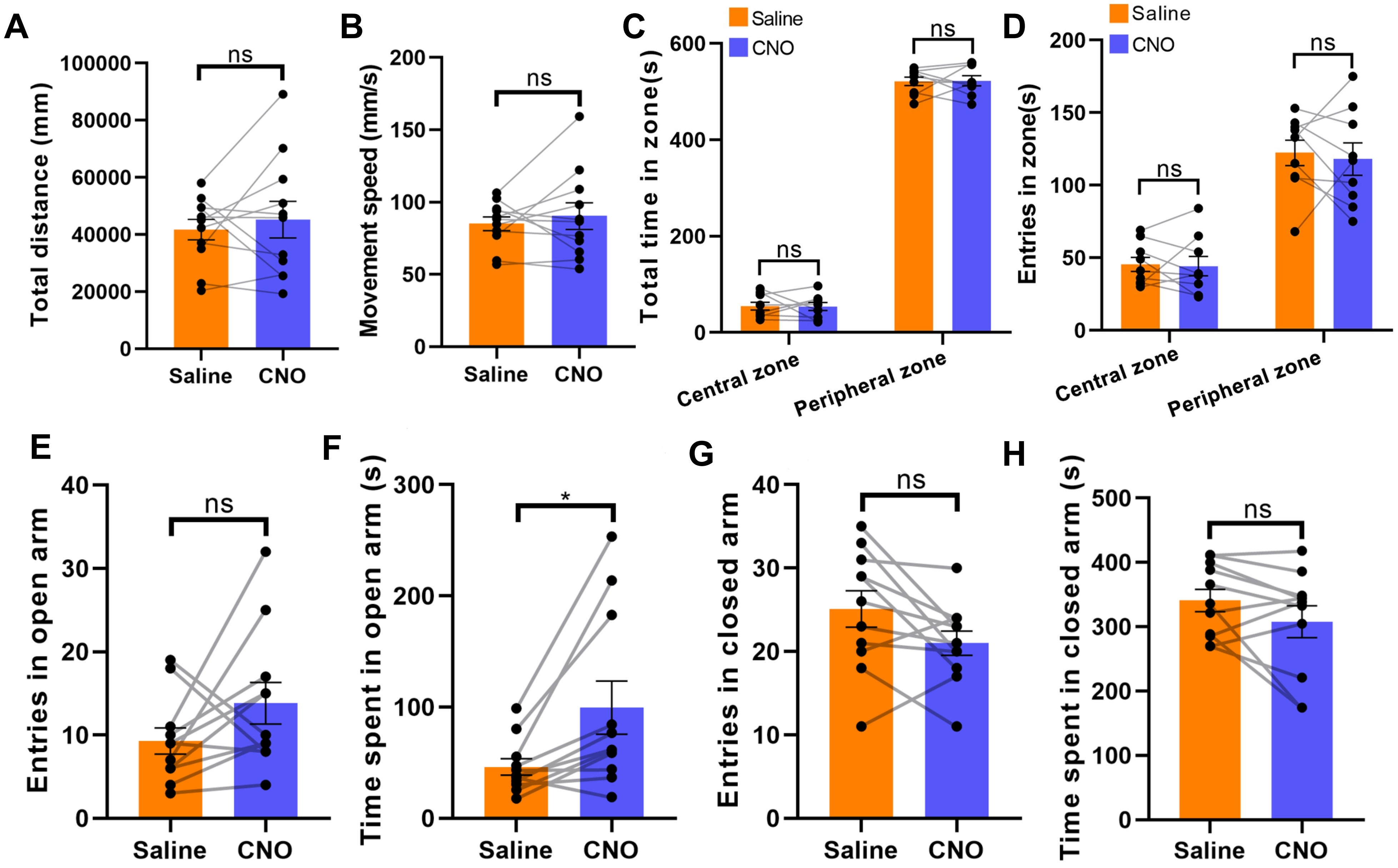

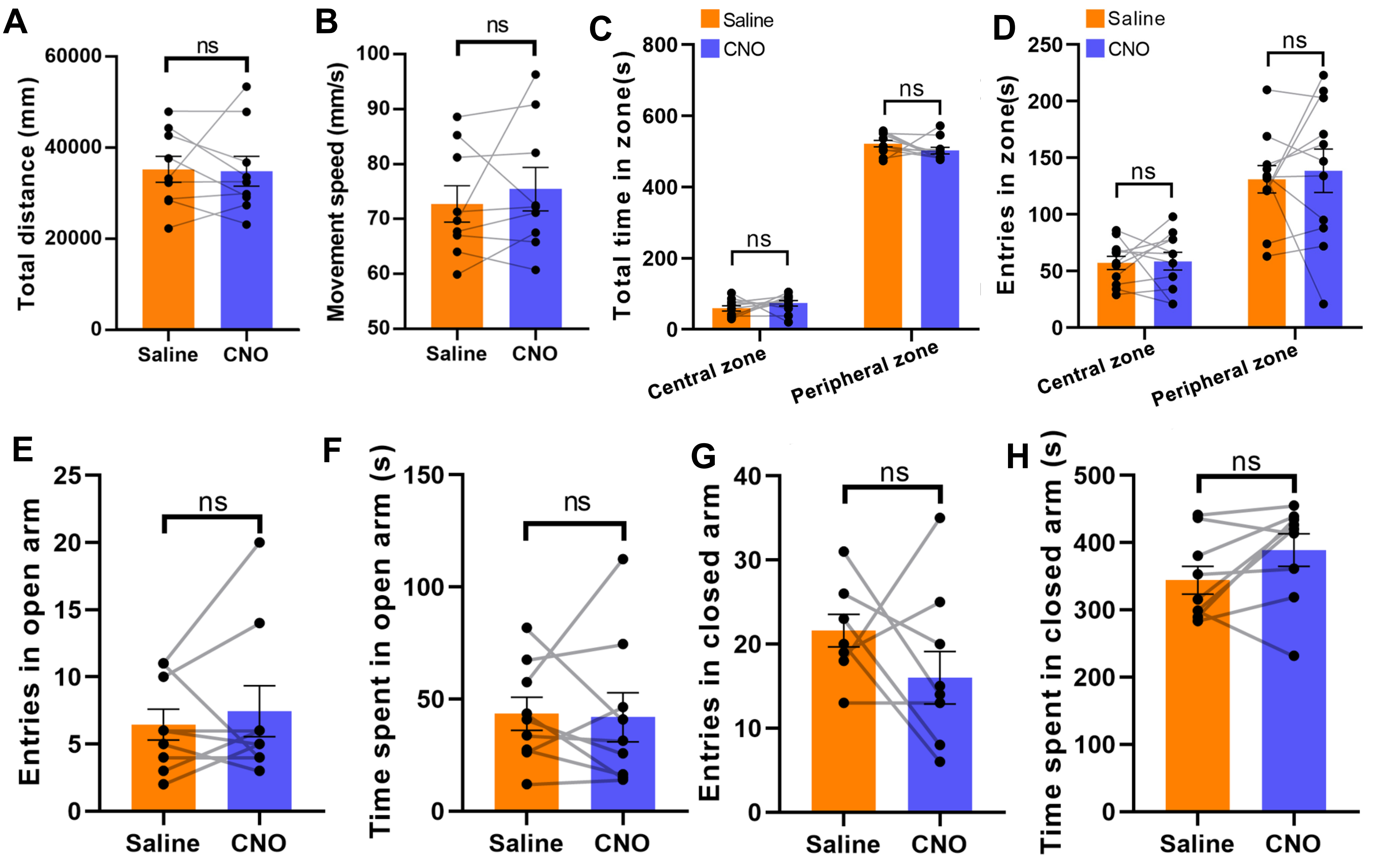

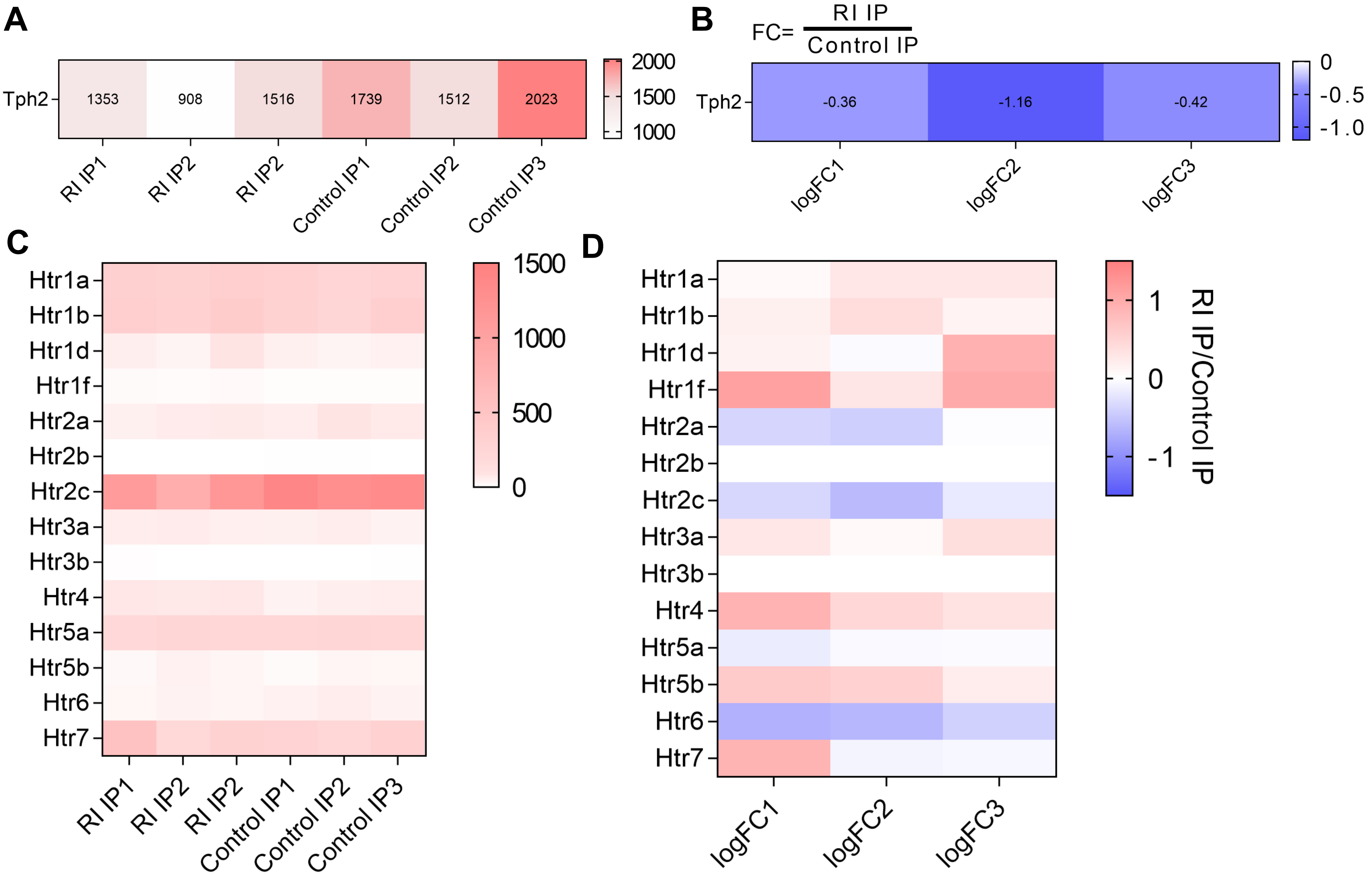
